# The embryonic DNA methylation program modulates the *cis-*regulatory landscape via CTCF antagonism

**DOI:** 10.1101/2023.11.16.567349

**Authors:** Ana Monteagudo-Sánchez, Julien Richard Albert, Margherita Scarpa, Daan Noordermeer, Maxim V.C. Greenberg

**Affiliations:** Université Paris Cité, CNRS, Institut Jacques Monod, F-75013 Paris, France; Université Paris-Saclay, CEA, CNRS, Institute for Integrative Biology of the Cell (I2BC), F-91998 Gif-sur-Yvette, France

## Abstract

During mammalian embryogenesis, both the 5-cytosine DNA methylation (5meC) landscape and three-dimensional (3D) chromatin architecture are profoundly remodeled during a process known as “epigenetic reprogramming.” An understudied aspect of epigenetic reprogramming is how the 5meC flux, *per se*, affects the 3D genome. This is pertinent given the 5meC-sensitivity of DNA binding for a key regulator of chromosome folding: CTCF. We profiled the CTCF binding landscape using a mouse embryonic stem cell (ESC) differentiation protocol that models the exit of naïve pluripotency, wherein global DNA methylation levels start low and increase to somatic levels within four days. We took advantage of the fact that mouse ESCs lacking DNA methylation machinery exhibit globally similar differentiation dynamics, thus allowing for dissection of more subtle effects of CTCF misregulation on gene expression. We carried this out by performing CTCF HiChIP in both wild-type and mutant conditions to assess aberrant CTCF-CTCF contacts in the absence of 5meC. We went on to assess the impact that misregulated CTCF binding has on *cis-*regulatory contacts by performing H3K27ac HiChIP, given that H3K27ac is enriched on active promoters and enhancers. Using DNA methylation epigenome editing, we were able to directly demonstrate that the DNA methyl-mark is able to impact CTCF binding. Finally, a detailed dissection of the imprinted *Zdbf2* gene showed how 5meC-antagonism of CTCF allows for proper gene regulation during differentiation. This work provides a comprehensive overview of how DNA methylation impacts the 3D genome in a relevant model for early embryonic events.

## Introduction

5-Cytosine DNA methylation (5meC) is a highly conserved epigenetic mark, generally associated with gene repression. In mammals, DNA methylation is typically found in the CpG dinucleotide context and approximately 80% of CpGs are methylated in somatic tissues. During the early stages of mammalian development following fertilization, most of the gametic 5meC is erased; subsequently, during implantation, the *de novo* methyltransferases DNA METHYLTRANSFERASE 3A and 3B (DNMT3A and DNMT3B) rapidly establish the embryonic DNA methylation landscape (Greenberg and Bourc’his 2019). This period is called the naïve-to-primed pluripotency transition, and occurs just prior to germ layer specification (Nichols et al. 2009). Changes in the histone modification patterns and in the transcriptional landscape are also substantial during this period (Argelaguet et al. 2019). Thus, it is presumed that the epigenome plays an integral role in preparing the cells within the embryo for lineage commitment.

In mammalian cell nuclei, the chromatin is organized in hierarchical structures that range from multi-megabase chromosome territories to more local cis-regulatory contacts (Lieberman-Aiden et al. 2009; Cremer and Cremer 2010; Rao et al. 2014; Bonev et al. 2017). Inside the territories, different chromatin compartments are defined by their transcriptional activity: the euchromatic “A” compartments that are typically transcriptionally active and the “B” compartments that are relatively transcriptionally repressed (Chen et al. 2018; Lieberman-Aiden et al. 2009). The compartments themselves are organized into Topologically Associating Domains (TADs)—“regulatory neighborhoods’’ that facilitate gene expression programs (Dixon et al. 2012; da Costa-Nunes and Noordermeer 2023; Nora et al. 2013). Within TADs, DNA loops can be formed, which are the smallest degree of organization, and can enable or insulate interactions between gene promoters and *cis*-regulatory elements such as enhancers (Tolhuis et al. 2002; Rao et al. 2014). The *cis* regulatory contacts differ substantially between cell types and they are crucial for determining proper cell identity (Schoenfelder and Fraser 2019; Zheng and Xie 2019). This hierarchical chromosome organization is dynamic and very important for several genomic processes, including transcription, gene regulation, replication and cell division (Monk 2015).

There are several architectural proteins involved in chromatin organization, CCCTC-BINDING FACTOR (CTCF) being one of the most well-characterized. CTCF is a zinc finger (ZF) protein that is highly conserved in mammals and that binds pervasively throughout the genome. It is known that CTCF plays a role together with the cohesin complex in the demarcation of TADs boundaries (de Wit et al. 2015; Li et al. 2020; Nora et al. 2017) and also has a role in transcription as it regulates loops between enhancers and promoters (Arzate-Mejía et al. 2018; Kubo et al. 2021). The absence of CTCF in mice is lethal in the early embryo, whereas heterozygous deletions of the protein present predisposition to cancer (Moore et al. 2012; Kemp et al. 2014), indicating that CTCF plays an essential role in development and cell identity. Genome profiling analysis of CTCF occupancy in human cells, obtained from different tissues, have shown considerable cell-type variability (Wang et al. 2012).

What are the mechanisms that dictate cell-specific CTCF binding patterns? Certainly, transcription factors play a role (Sahu et al. 2022; Kreibich et al. 2023), as well as chromatin modifying complexes (Kaaij et al. 2019). However, a compelling mechanism is DNA methylation itself, given that substantial variability of CTCF binding patterns between cell types can be linked to 5meC status in the binding site (Wang et al. 2012). Biochemical analyses have confirmed that the presence of 5meC at certain cytosines within the CTCF binding motif can significantly impair CTCF-DNA interactions (Hashimoto et al. 2017). While DNA methylation does not appear to play an important role in either TAD or compartment establishment (Hassan-Zadeh et al. 2017; Nothjunge et al. 2017; Jiang et al. 2020a), 5meC indeed has an effect on relatively short *cis* regulatory contacts in some contexts. *In vivo* the antagonistic relation between CTCF binding and 5meC has been well-documented at genomic imprints. For example, the paternally methylated *H19-Igf2* imprinting control region (ICR) repels CTCF binding, allowing for interactions between enhancers at this locus and the *Igf2* promoter leading to expression. Conversely, CTCF binds the unmethylated maternal allele, insulating the activation of the *Igf2* promoter from its enhancers, which in turn allows the expression of the *H19* long non-coding RNA (Hark et al. 2000; Bell and Felsenfeld 2000; Llères et al. 2019). The antagonism between CTCF binding and 5meC has been also observed in tumors (Fang et al. 2020); in IDH mutant gliomas the hypermethylation of a CTCF binding site causes a reduction in CTCF binding that results in the expression of a glioma oncogene (Flavahan et al. 2016; Rahme et al. 2023).

In this study we set out to determine how the dramatic embryonic DNA methylation program impacts three-dimensional chromatin architecture and underlying gene regulation in a dynamic system. We employed a mouse embryonic stem cell (ESC) differentiation approach that recapitulates the embryonic *de novo* DNA methylation program: naïve ESCs cultured in serum-free media, which are characterized by low levels of DNA methylation (Blaschke et al. 2013), were differentiated to Epiblast-like cells (EpiLCs) inducing the transition towards primed pluripotency (Guo et al. 2009). In parallel, we employed a *Dnmt1; Dnmt3a; Dnmt3b* triple knockout (TKO) cell line, which despite completely lacking DNA methylation (Tsumura et al. 2006), is able to adopt a primed-like state during EpiLC differentiation (Greenberg et al. 2017, 2019; Schulz et al. 2022; Richard Albert et al. 2023). We profiled CTCF binding changes in ESCs and EpiLCs in the presence and absence of 5meC, showing that ∼1% CTCF binding sites are enriched in TKO EpiLCs, relative to WT. Previous chromosome conformation studies using DNA methylation mutants were not able to detect architectural differences at finer scales, therefore likely missed many short range *cis*-regulatory interactions (Jiang et al. 2020a; Nothjunge et al. 2017). Hence, we utilized HiChIP (Mumbach et al. 2016), which captures both short and long-range interactions, and allowed us to assess either chromatin loops by enriching for CTCF-bound loci, or enhancer-promoter contacts by enriching for histone H3 lysine 27 acetylation (H3K27ac)-marked regions. We could then determine how misregulated CTCF affects the *cis*-regulatory landscape. We functionally demonstrated that 5meC negatively impacts CTCF binding at multiple loci by implementing epigenome editing. Finally, we carried out fine grained genetic experiments to show how 5meC influences CTCF-mediated gene regulation at the imprinted *Zdbf2* locus. In sum, our study provides a comprehensive view of how the embryonic DNA methylation program contributes to chromatin folding as a means to control gene expression.

## Results

### DNA methylation perturbs CTCF binding at minority of sites

In naïve mouse ESCs, the *de novo* and maintenance DNA methylation is impaired, while active DNA demethylation is stimulated, leading to extremely low 5meC levels: <10% of all CpGs are methylated, mainly localized to transposable elements (Leitch et al. 2013; Hackett and Surani 2014; Marks et al. 2012; von Meyenn et al. 2016; Walter et al. 2016) (Figure 1A,B). To achieve this state, we cultured ESCs in serum-free media, supplemented with MEK and GSK3β inhibitors plus vitamin C (2i+vitC) (Blaschke et al. 2013). Given the global DNA hypomethylation, perhaps unsurprisingly the transcriptional landscape of WT naïve ESCs is highly similar with that of TKO ESCs cultured in the same conditions (Schulz et al. 2022). We went on to profile the CTCF binding in both WT and TKO ESCs by Cleavage Under Targets and Release Using Nuclease (CUT&RUN) (Skene and Henikoff 2017). Consistent with the DNA methylation and transcriptomic data, the CTCF binding patterns are coherent between WT and mutant conditions (Supplementary Figure S1A,B).

**Figure 1.**
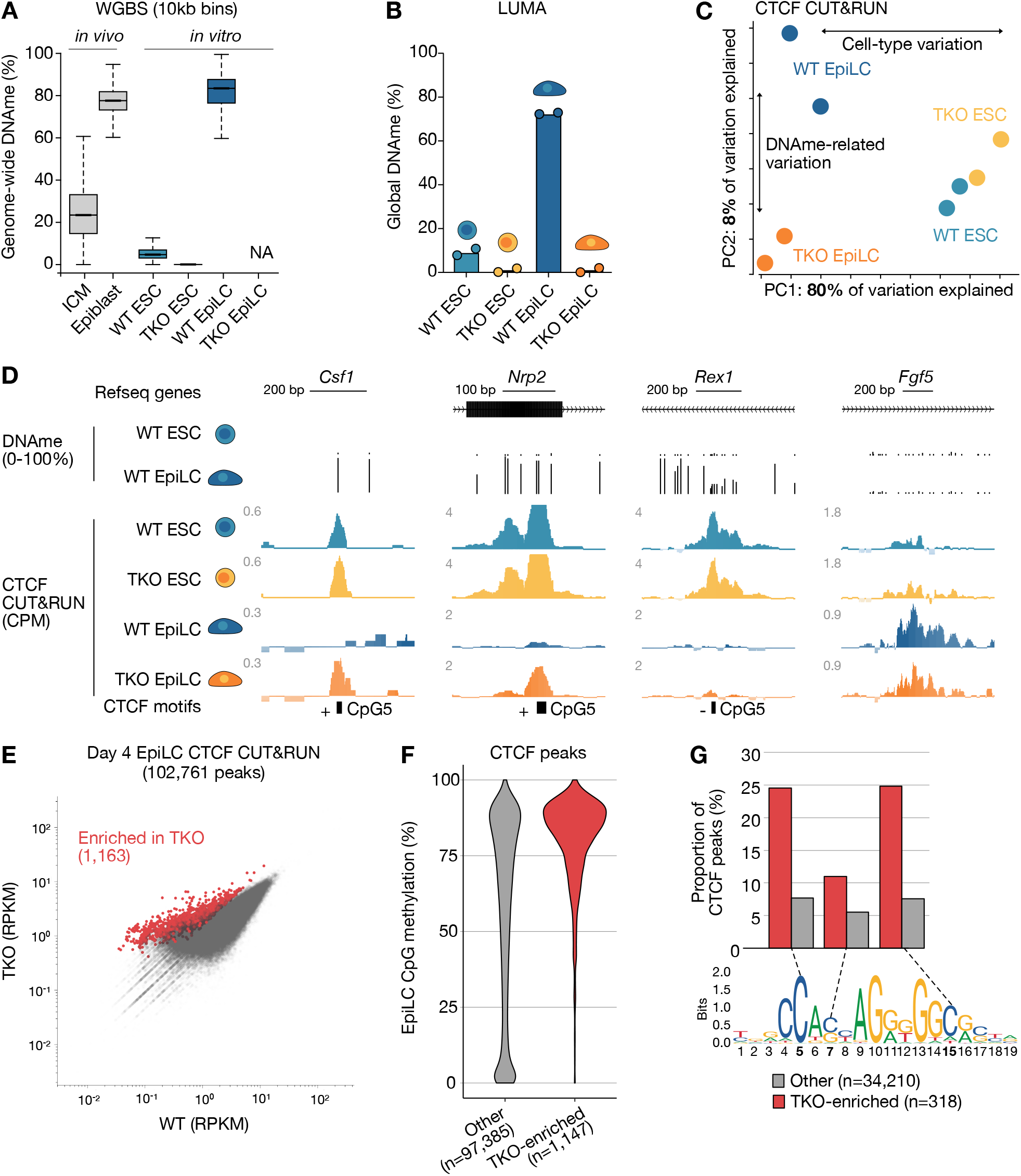
CTCF exhibits potential 5meC-sensitivity at a substantial number of loci. **A.** Distribution of average 5meC levels over 10kb bins (n=273,121) in E3.5 inner cell mass cells (ICM), the E7.5 epiblast, *Dnmt* WT (WT) and *Dnmt* triple KO (TKO) embryonic stem cells (ESCs) and Epiblast-like cells (EpiLCs). Data from Wang et al. 2014 (*in vivo*), Richard Albert et al. 2023 (WT ESC & EpiLC) and Domcke et al. 2015 (TKO ESC). Boxplots represent the median (line inside the box), where 50% of the data are distributed (the box), and whiskers denote the values lying within1.5 times the interquartile range. **B**. Luminometric methylation assay (LUMA) depicting global 5meC levels in WT ESCs grown in serum and 2i+vitC, TKO ESCs grown in 2i+vitC, and WT and TKO EpiLCs. Data are represented as the mean of two replicates, which are included as individual data points. **C**. Principal component analysis (PCA) plot of CTCF CUT&RUN data in WT and TKO ESCs and EpiLCs. Individual replicates and the percent variation explained by each principal component are shown. **D**. UCSC genome browser screenshots of putative 5meC-sensitive CTCF binding sites at the *Csf1* and *Nrp2* loci. DNA methylation levels in WT ESCs and EpiLCs and the location of Refseq genes and CTCF binding motifs are included. The strand (+/-) and position (5, 7 or 15) of the methylated CpG is indicated. Cell-type specific CTCF binding at the naïve pluripotency marker gene *Rex1* and the primed pluripotency marker gene *Fgf5* are shown for comparison. *Csf1*: chr3:107,728,899-107,729,432, *Nrp2*: chr1:62,738,262-62,738,551, *Rex1*: chr8:43,305,548-43,306,216, *Fgf5*: chr5:98,255,482-98,256,478. **E**. 2D scatterplot showing CTCF peak enrichment levels (RPKM) in WT and TKO EpiLCs. Statistically enriched peaks in TKO (linear modeling with Limma, FC>2, t-test adjusted p value <0.05) are highlighted in red. **F.** Violin plot of the distribution of EpiLC DNA methylation levels within CTCF peaks. Peaks are categorized as in **E**. **G.** Bar plot showing the proportion of CTCF peaks that overlap a canonical CTCF binding motif with a CpG at position 5, 7 or 15. The CTCF sequence motif is included (JASPAR MA0139.1).

Next we differentiated WT and TKO ESCs to EpiLCs. Within four days, the WT genome is highly methylated (∼80% methylated CpGs), whereas the TKO EpiLC genome remains completely unmethylated (Greenberg et al. 2019) (Figure 1A, B). Nevertheless, even in the absence of DNA methylation, we and others have previously demonstrated TKO can not only exit naïve pluripotency, but can do so with similar differentiation kinetics as WT (Greenberg et al. 2017; Hassan-Zadeh et al. 2017; Schulz et al. 2022). We reasoned then that EpiLC differentiation would provide a dynamic system in which we could directly compare cell type-specific versus DNA methylation-mediated CTCF regulation. Therefore, as with ESCs, we performed CTCF CUT&RUN in WT and mutant day 4 (D4) of differentiation EpiLCs. The CTCF binding landscape in WT and TKO EpiLCs globally resembled each other more than the TKO EpiLCs resembled naïve ESCs (Figure 1C; Supplementary Figure S1A). This suggests that the cell type plays a more important role in determining CTCF occupancy than DNA methylation, *per se*. Although it is worth noting that there are more differences between WT and TKO EpiLCs than there are between WT and TKO ESCs (Figure 1C; Supplementary Figure S1A,B).

Nevertheless, we were able to determine a substantial number of CTCF peaks that were enriched specifically in TKO EpiLCs: 1,163 out of 102,761 total peaks (FC≥2, adjusted p<0.05) (Figure 1D,E; Supplementary Figure S1B). A number of data points indicate that DNA methylation is antagonizing CTCF binding at these elements. Firstly, TKO-specific peaks were depleted at promoter regions, which are generally DNA methylation-free (Supplementary Figure S1C). Secondly, using whole genome bisulfite (WGBS) data (Richard Albert et al. 2023), we could observe that the vast majority of TKO-enriched sites gain DNA methylation in WT (Figure 1F; Supplementary Figure S1D). This stood in contrast with all other peaks, which exhibited much less bias for DNA methylation gain, and harbored a substantial number of sites that remained unmethylated. Thirdly, we examined in detail the DNA methylation state of CpGs within the CTCF binding motif at the TKO-specific sites. The CTCF core binding motif may contain a number of CpG sites that have been demonstrated to impact CTCF-DNA interactions when methylated, with position five in the JASPAR motif (Figure 1G) exhibiting the most substantial effect in biochemical studies (Maurano et al. 2015; Hashimoto et al. 2017; Kreibich et al. 2023). Of the 1,163 TKO-enriched sites, we were able to determine 318 wherein we could discern a CTCF motif with DNA methylation information at least one pertinent CpG (Figure 1G). This may indicate that DNA methylation in the local chromatin environment, perhaps via interaction with methyl-sensitive DNA binding proteins, may play a substantial role in shaping the CTCF binding landscape (Wiehle et al. 2019). Nevertheless, we focused on CTCF binding sites containing CpGs. Consistent with the overall DNA methylation pattern (Figure 1F), all three CpG sites showed an enrichment of DNA methylation compared to all other peaks (Figure 1G). While the overall percentage of TKO-specific peaks is only ∼1% of the total number of binding sites, there still remains a fairly considerable number of sites where DNA methylation can influence not only CTCF binding, but potentially *cis* regulatory gene control.

### 3D genome architecture is globally preserved in DNA methylation-deficient pluripotent cells

HiChIP is a variation of Hi-C in which cross-linked chromatin is immunoprecipitated for a chromatin-associated factor or modification prior to sequencing (Mumbach et al. 2016). We performed HiChIP of CTCF in WT and TKO ESCs and EpiLCs in order to enrich for CTCF-anchored contacts (Supplementary Table S1). The HiChIP data is versatile in that both CTCF occupancy, as well as genomic contact information data is generated. Consistent with the CTCF CUT&RUN profiles, the HiChIP datasets are grouped more closely by cell type as opposed to genotype (Figure 2A, Supplementary Figure S2A). Moreover, the data indicated that CTCF enrichment over TAD borders and A/B compartment organization was largely unperturbed in the absence of DNA methylation (Supplementary Figure S2B-D). From merging the data by cell type, we were able to determine 126,481 total CTCF peaks in ESCs compared with 135,165 in EpiLCs, thus a very modest 7% increase. We next analyzed the number of significant contacts from the HiChIP data, focusing on those contacts that link CTCF-bound sites together. The merged data from WT samples revealed 201,998 total contacts. Using stringent parameters, we performed differential analyses and uncovered 911 ESC-specific and 1,726 EpiLC-specific loops that met our significance thresholds (FDR≤0.05, logFC≥1) (Figure 2B). It is worth noting that previous studies have also reported an increase in CTCF-CTCF contacts during ESC differentiation, which may signify cell type-specific gene regulatory programs becoming cemented (Bonev et al. 2017).

**Figure 2.**
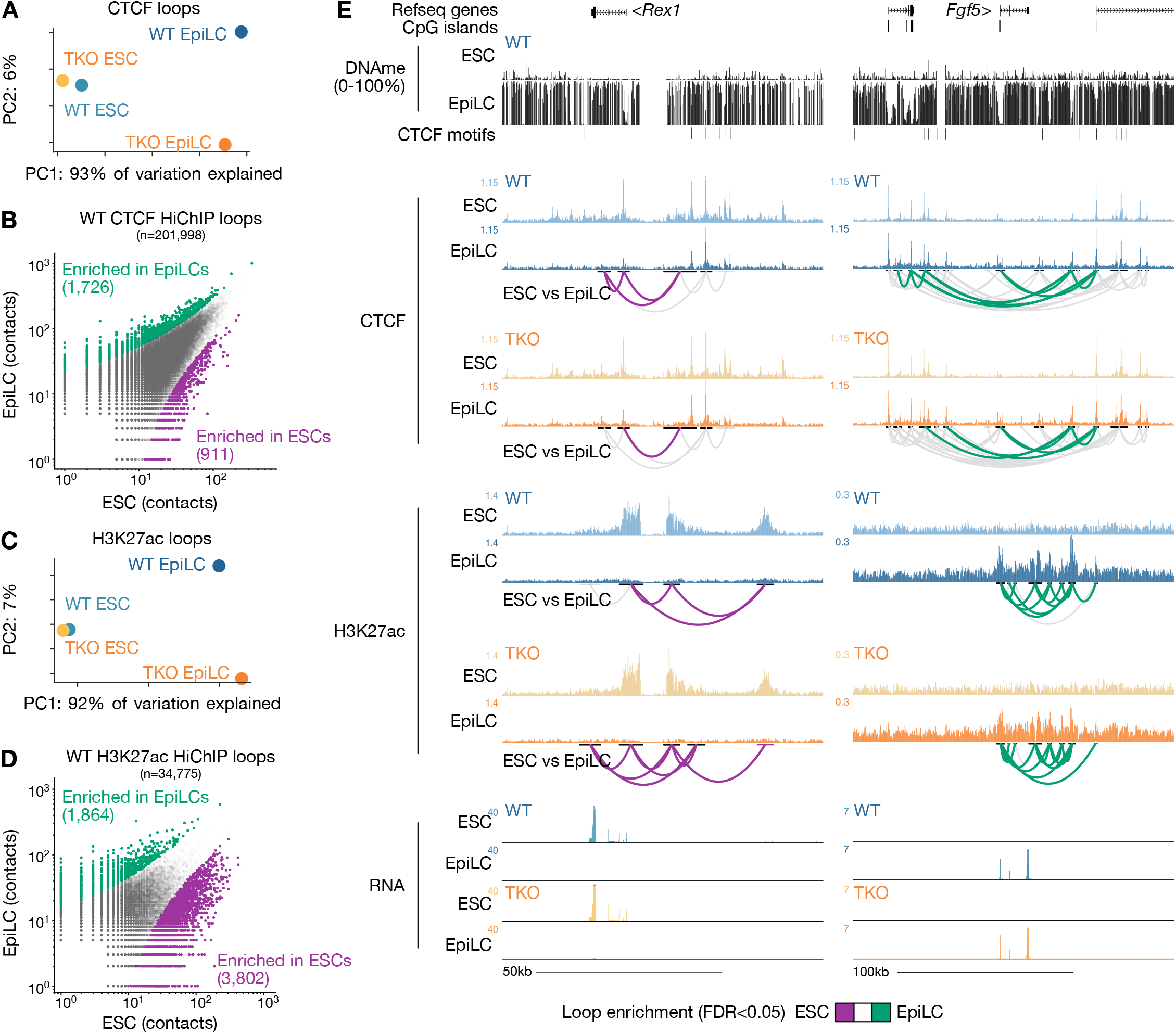
Chromatin architecture is remodeled during EpiLC differentiation in WT and TKO backgrounds. **A.** PCA plot showing the variation in CTCF contacts between WT and TKO ESCs and EpiLCs. **B.** 2D scatterplot of CTCF contacts between WT ESCs and EpiLCs. All significant loops are shown in gray. Loops with enriched contacts (>4 reads, FDR<0.05) in EpiLCs are highlighted in green and ESC-enriched loops are highlighted in purple. **C**. PCA plot showing the variation in 3D H3K27ac interaction contacts between WT and TKO ESCs and EpiLCs. **D.** 2D scatterplot of H3K27ac contacts between WT ESCs and EpiLCs as in **B**. **E.** UCSC genome browser screenshots of the naïve pluripotency marker gene *Rex1* and the formative pluripotency marker gene *Fgf5* loci. Note the ESC-specific CTCF and H3K27ac binding, CTCF and H3K27ac looping, and gene expression of *Rex1* in WT and TKO backgrounds. EpiLC-specific binding, loop contacts and expression of *Fgf5* is included for comparison. *Rex1*: chr8:43,264,884-43,373,313, *Fgf5*: chr5:98,139,881-98,390,116.

Given the absence of an effect on large chromatin structures, we reasoned that CTCF misregulation may rather impact relatively shorter *cis* regulatory contacts (Ren et al. 2017). Thus, we performed H3K27ac HiChIP in WT and TKO ESCs and EpiLCs in order to establish the “enhancer connectome” in each of these conditions (Mumbach et al. 2017). Keeping in line with the CTCF and transcriptome data, the H3K27ac landscape clusters by cell type much more strongly than by genotype (Figure 2C, Supplementary Figure S2E). We were able to identify 3,802 ESC-specific H3K27ac contacts, and 1,864 in EpiLCs (Figure 2D). This can be readily observed at marker genes for ESCs and EpiLCs, respectively, which exhibited dramatic changes in their H3K27ac-enriched enhancer-promoter contacts independently of the DNA methylation state (Figure 2E). The global preservation of chromatin architecture in TKO EpiLCs strongly bolsters our previous findings that DNA methylation is dispensable for exiting naïve pluripotency (Schulz et al. 2022). However, we were curious to pursue whether we could discover a class of genes that are sensitive to CTCF misregulation in the 5meC mutant, even if the impact on the overall EpiLC state may be more nuanced.

In line with the fact that both WT and TKO ESCs exhibit low/absent levels of 5meC, we only observed two differential loops in this cell type (Figure 3A). More saliently, CTCF-anchored loops enriched in TKO EpiLCs relative to WT—where 5meC levels are very high—may indicate DNA methylation sensitivity. Using the same analysis, we uncovered 43 differential loops in the DNA methylation mutant, using stringent thresholding parameters (Figure 3A). Notably, only three loops were enriched in the WT EpiLCs. Analyzing a larger region around each anchor (+/- 1750bp) identifies an additional 75 TKO-specific CTCF loops, and 15 in WT. Thus, by combining both analyses, we defined a final list of 118 differential CTCF contacts, with a noticeable enrichment of contacts in TKO EpiLCs. Consistently, CTCF peaks from the TKO EpiLC HiChIP data were predominately DNA methylated in WT (Supplementary Figure S3A,B). It is also worth emphasizing that CpG at position 5 was more enriched than other CpGs in the CTCF binding motif in the TKO EpiLC HiChIP data, suggesting that this is the most deterministic base for 5meC-mediated antagonism (Supplementary Figure S3C).

**Figure 3.**
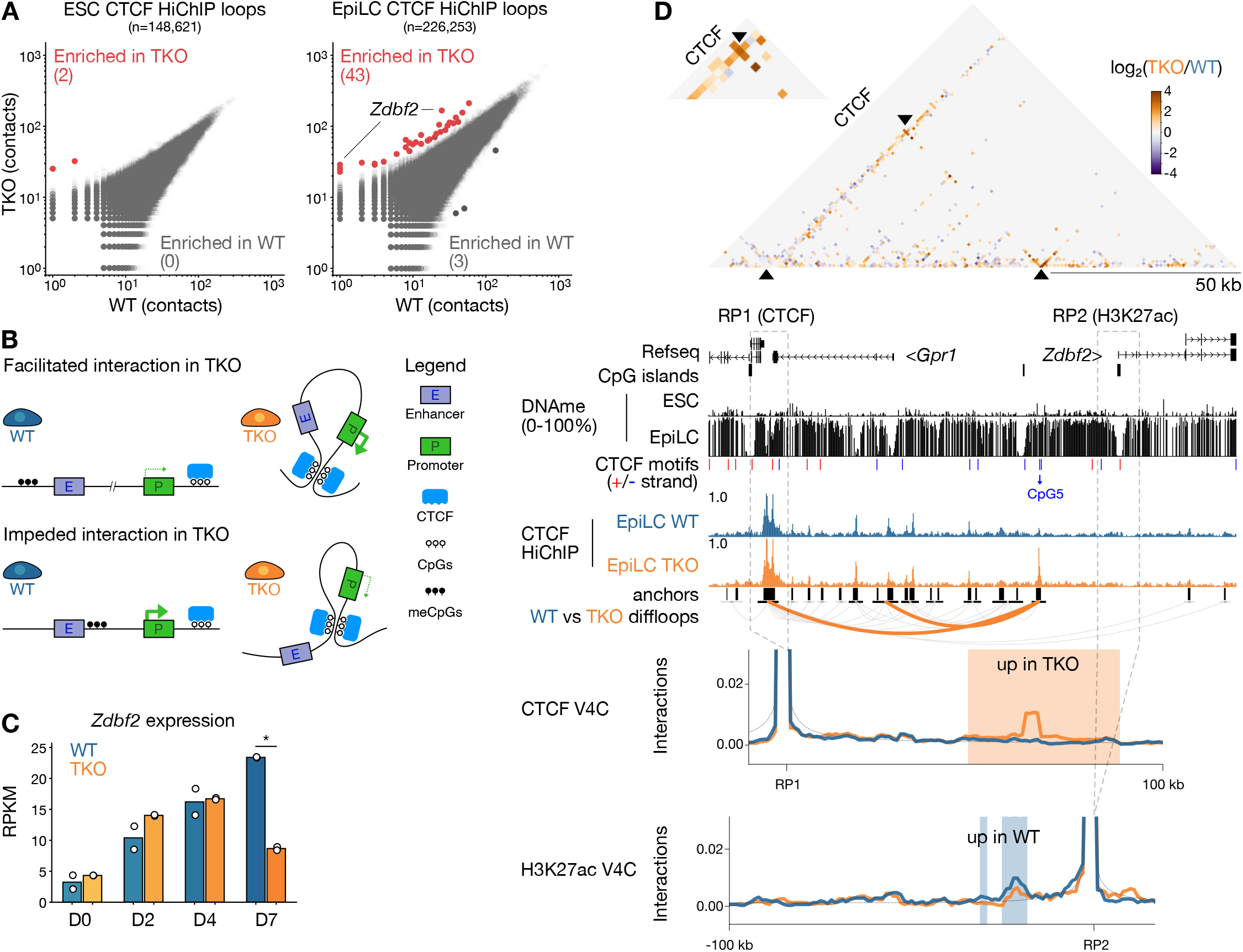
A subset of CTCF loops and promoter-enhancer contacts are disrupted in 5meC-deficient EpiLCs. **A.** 2D scatterplots of CTCF loop contacts in WT versus TKO ESCs (left) and EpiLCs (right). Loops with significantly enriched contacts in TKO cells (log₂FC>1, FDR<0.05) are highlighted in red. **B.** Schema showing two potential scenarios resulting from 5meC-sensitive CTCF binding and altered promoter-enhancer contact formation and nearby gene transcription. **C.** Bar plot of *Zdbf2* expression levels (RPKM) in a time course assay of ESC-to-EpiLC differentiation in WT and TKO cells. Bars represent the mean, and replicates are shown as dots. D: days post-AF treatment. Statistically significant changes in gene expression between WT and TKO (Linear modeling with Limma, FC>1, t-test adjusted p value <0.05) are indicated by asterisks. **D.** Contact matrix (top), genome browser screenshot (middle) and virtual 4C (bottom) plots of the *Zdbf2* locus. Top: differential CTCF contacts between WT and TKO EpiLCs are displayed, where each pixel represents a 1 kb bin. The TKO-enriched contacts between the TAD border and the putative DNAme-sensitive CTCF binding site is magnified in the inset. Reference points 1 (CTCF) and 2 (H3K27ac) for the virtual 4C plots (bottom) are indicated by a dashed box. Middle: browser screenshot showing CpG methylation, CTCF enrichment levels and loops. Refseq genes, CpG islands, CTCF motifs (positive strand in red, negative strand in blue) are included. The CTCF motif with a CpG at position 5 that underlies a putative 5meC-sensitive CTCF peak is highlighted in blue. Bottom: Virtual 4C plots of reference points 1 (CTCF) and 2 (H3K27ac) showing interaction frequencies between the reference point and adjacent area. The background model is shown as a dotted gray line. Statistically enriched contacts (chi-squared test, alpha<0.25) are highlighted in blue (high in WT EpiLC) or orange (high in TKO EpiLC). Coordinates: chr1:63,165,972-63,304,123.

### TKO EpiLC-specific CTCF loops are correlated with gene misregulation at discrete loci

CTCF-mediated chromosome folding can ensure enhancer-promoter contacts allowing for proper gene expression, and at the same time insulate promoters from aberrant enhancer interactions (Kim et al. 2015) (Figure 3B). Therefore, we set out to determine if the *de novo* DNA methylation program can exert an effect on CTCF-dependent gene control. Our strategy was to systematically assess the 118 differential loops that are enriched in TKO EpiLCs, and determine if H3K27ac contacts and gene expression were impacted. We identified 163 genes that indicate that DNA methylation could influence gene expression via CTCF antagonism based on their proximity to misregulated CTCF contacts.

We culled this list to four compelling loci for further analysis: *Csf1*, *Mob3b, Nrp2* and *Zdbf2*. The *Csf1, Mob3b* and *Nrp2* genes either contain or are adjacent to CTCF binding sites containing a CpG at position 5, and CTCF was enriched at these sites in TKO EpiLCs but not WT (Supplementary Figure S4, S5, and S6). Importantly, in all cases the TKO EpiLC-specific binding was associated with the formation of differential loop(s) (FDR<0.05, FC≥2). Finally, these three genes were upregulated TKO EpiLCs (Supplementary Figure S7A). At least in the case of *Csf1* and *Mob3b*, virtual 4C suggested that the TKO EpiLC-specific loop was also associated with increased interactions between H3K27ac-enriched regions (Supplementary Figure S4 and S5). In other words, the aberrant CTCF-CTCF looping could be facilitating enhancer-promoter contacts, leading to upregulation (Figure 3B).

Finally, from our analyses, the most significant differential CTCF loop that was enriched in TKO EpiLCs was found at the imprinted *Zdbf2* locus. However, as opposed to the previous examples, the presence of the aberrant loop was correlated with decreased *Zdbf2* expression (Figure 3C). While we uncovered this loop through an agnostic approach, incidentally this is a locus that we have previously characterized. During ESC to EpiLC differentiation, DNA methylation upstream of the *Zdbf2* promoter is required to antagonize polycomb repressive complex 2 (PRC2)-mediated silencing in order to allow proper gene activation (Duffié et al. 2014; Greenberg et al. 2017). We also showed that four enhancers upstream of the *Zdbf2* promoter are crucial for its activity (Greenberg et al. 2019). The physiological consequences of embryonic *Zdbf2* regulation are life-long: in mouse embryos where DNA methylation is not deposited upstream of the *Zdbf2* promoter, the gene remains constitutively polycomb-repressed, leading decreased appetite, smaller size, and lower survivability in affected pups with respect to their WT littermates (Glaser et al. 2022; Greenberg et al. 2017). As such, *Zdbf2* has emerged as a valuable locus to study the long-lasting effects of epigenetic reprogramming.

The differential loop at the *Zdbf2* locus is anchored in a CTCF binding site that sits between the *Zdbf2* promoter and the aforementioned four enhancers. In WT EpiLCs, CTCF binding was depleted, which was correlated with a gain of 5meC at position 5 in its binding site (Figure 3D). In TKO EpiLCs, where CTCF binding was maintained, our H3K27ac HiChIP data revealed less interactions between the *Zdbf2* promoter with upstream enhancers in the TKO EpiLCs compared with WT (Figure 3D). Thus, we reasoned that in ESCs, the CTCF binding could help insulate *Zdbf2* from precocious activation; this insulation is maintained in the DNA methylation mutant, helping to explain the persistent repression (Figure 3B).

### Epigenome editing confirms DNA methylation-CTCF antagonism

While globally the WT and TKO EpiLCs are transcriptionally similar, there are a substantial number of misregulated genes in the DNA methylation mutant (Schulz et al. 2022). Thus, it is possible that the gene misregulation we have described may be indirect of CTCF-mediated action. To formally demonstrate that DNA methylation*, per se*, affects CTCF binding and downstream regulatory defects, we targeted locus specific DNA demethylation using the CRISPR/Cas9 SunTag system. Briefly, catalytically inactive Cas9 (dCas) fused to five GCN4 epitopes (SunTag) recruits the TET1 catalytic domain fused to GFP and a single chain variable fragment (scFv) that recognizes the SunTag (Morita et al. 2016). We took advantage of a piggyBac transgenesis-compatible plasmid where all components are expressed as a single transcript driven by a constitutive promoter, and the translated peptide contains the P2A self-cleavable peptide sequence between the Cas9-SunTag and GFP-scFv-TET1 (Supplementary Figure S7B) (Horii et al. 2022; Richard Albert et al. 2023). After selecting for GFP positive cells, we used piggyBac-mediated transgenesis to stably integrate single guide RNAs (sgRNAs) that target the epigenome editing machinery to the respective CTCF binding sites. Following selection of sgRNA integration, we differentiated the dCas9-SunTag/TET and control lines to EpiLCs for four days. In all cases, we would expect that persistent DNA demethylation would lead to increased CTCF binding. However, depending on the mode of regulation, we would expect either increased expression (eg, *Csf1*, *Mob3b* and *Nrp2*) or repression (eg, *Zdbf2*) (Figure 4A).

**Figure 4.**
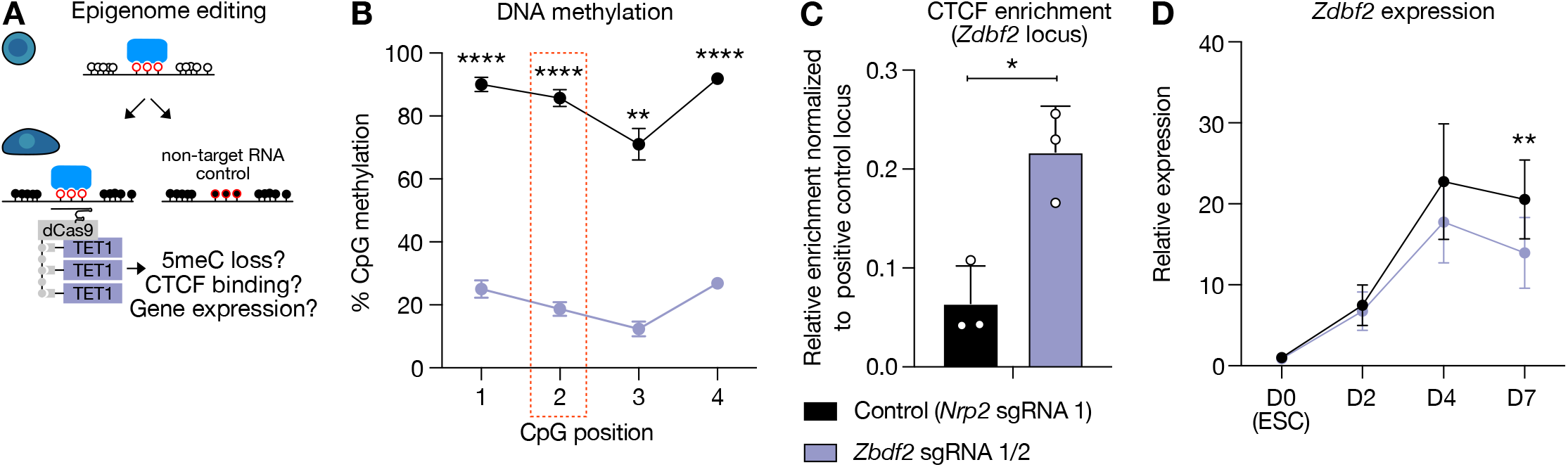
Precision cytosine demethylation at the *Zdbf2* locus results in increased CTCF binding and failure to completely activate gene expression. A. Schema depicting the site-directed 5meC erasure strategy. A catalytically inactive Cas9 (dCas9, gray rectangle) fused to a SunTag (five gray circles) is recruited to target chromatin by one or several short guide RNAs and in turn recruits the catalytic domain of TET1 (purple) via scFv interactions with the SunTag. Cells are differentiated for 4 days (left) and assessed for 5meC levels, CTCF binding and nearby gene expression compared to control cells expressing non-target RNA (right). **B.** Bisulphite-pyrosequencing results of cells expressing non-target sgRNA (black) and cells expressing *Zdbf2* CTCF site target sgRNAs (purple). The position of each CpG within the amplicon is indicated, and the CpG corresponding to CpG5 in the canonical CTCF binding motif is highlighted by the red box. Data are shown as mean ± standard error for three replicates. **C.** ChIP-qPCR results of the same cells as in **B**. Data are shown as mean ± standard error for three replicates represented by unfilled circles. **D.** RT-qPCR results of the same cells as in **B** over a time course of 7 days (D) of EpiLC differentiation. Expression of each replicate was normalized to two housekeeping genes (*Rrm2* & *Rplp0*), and then to WT ESCs. Data are shown as mean ± standard error for three replicates. p-values were calculated by one-tailed paired t-test assuming equal variance: *p<0.05, **p<0.01, **p<0.001, ****p<0.0001.

Indeed, we observed robust targeted DNA demethylation for each candidate locus compared to control (Figure 4B, Supplementary Figure S7C). Validating our prediction, reduced 5meC was associated with increased CTCF binding in every case (Figure 4C, Supplementary Figure S8A,B). We next assessed if local gene expression was altered when CTCF binding was increased. In the case of *Csf1*, *Mob3b* and *Nrp2*, we did not observe changes in expression that were consistent with our model (Supplementary Figure S8C). This could be due to the fact that the increased expression of these genes in the *Dnmt* TKO was due to secondary effects unrelated to misregulated CTCF. Alternatively, the lack of an effect could be because dCas9-SunTag/TET-mediated DNA demethylation was not complete, CTCF binding was only slightly altered, and a threshold was not attained that would impact transcriptional processes. It should also be noted that *Mob3b* expression actually decreased in the presence of the epigenome editing machinery, even in globally hypomethylated ESCs (Supplementary Figure S8C). This may indicate transcriptional interference by the dCas9-SunTag complex bound in the body of the *Mob3b* gene, which would confound our results. However, fitting with our model, the enriched CTCF at the epigenome-edited *Zdbf2* locus led to a significant decrease in *Zdbf2* expression (Figure 4D).

### CTCF and Polycomb both coordinate repression of Zdbf2 in the hypomethylated state

We were intrigued by our dCas9-SunTag/TET1 editing results at *Zdbf2*, and were motivated to perform further genetic tests to substantiate our model of CTCF insulating the four enhancers from contacting the *Zdbf2* promoter (Figure 5A,B). To do this, we generated a homozygous deletion mutant of the CTCF binding site in WT ESCs (Figure 5A, Supplementary Figure S9A). In ESCs lacking the CTCF binding site, indeed we observed a minor increase in *Zdbf2* expression (Figure 5C). As mentioned, *Zdbf2* is polycomb repressed in ESCs (Figure 5A,B), and we previously demonstrated that addition of a PRC2 inhibitor to ESC culture media leads to mild de-repression (Greenberg et al. 2019). We performed the same experiment here, and recapitulated the mild effect observed in the WT background (Figure 5C, Supplementary Figure S9B). Strikingly, we observed a substantial upregulation (∼17 fold) when we added the PRC2 inhibitor to cells lacking the CTCF binding site (Figure 5C). These data strongly suggest that Polycomb and CTCF synergistically cooperate to maintain *Zdbf2* repression in the hypomethylated state—H3K27me3 is enriched TKO EpiLCs as well (Greenberg et al. 2017; Richard Albert et al. 2023)—and the *de novo* DNA methylation program is required to release both of these means of control (Figure 5A-C). Finally, consistent with our prediction, in EpiLCs when the DNA methylation levels are high and CTCF is no longer bound, the deletion of the CTCF binding site did not lead to an effect on *Zdbf2* expression (Supplementary Figure S9C).

**Figure 5.**
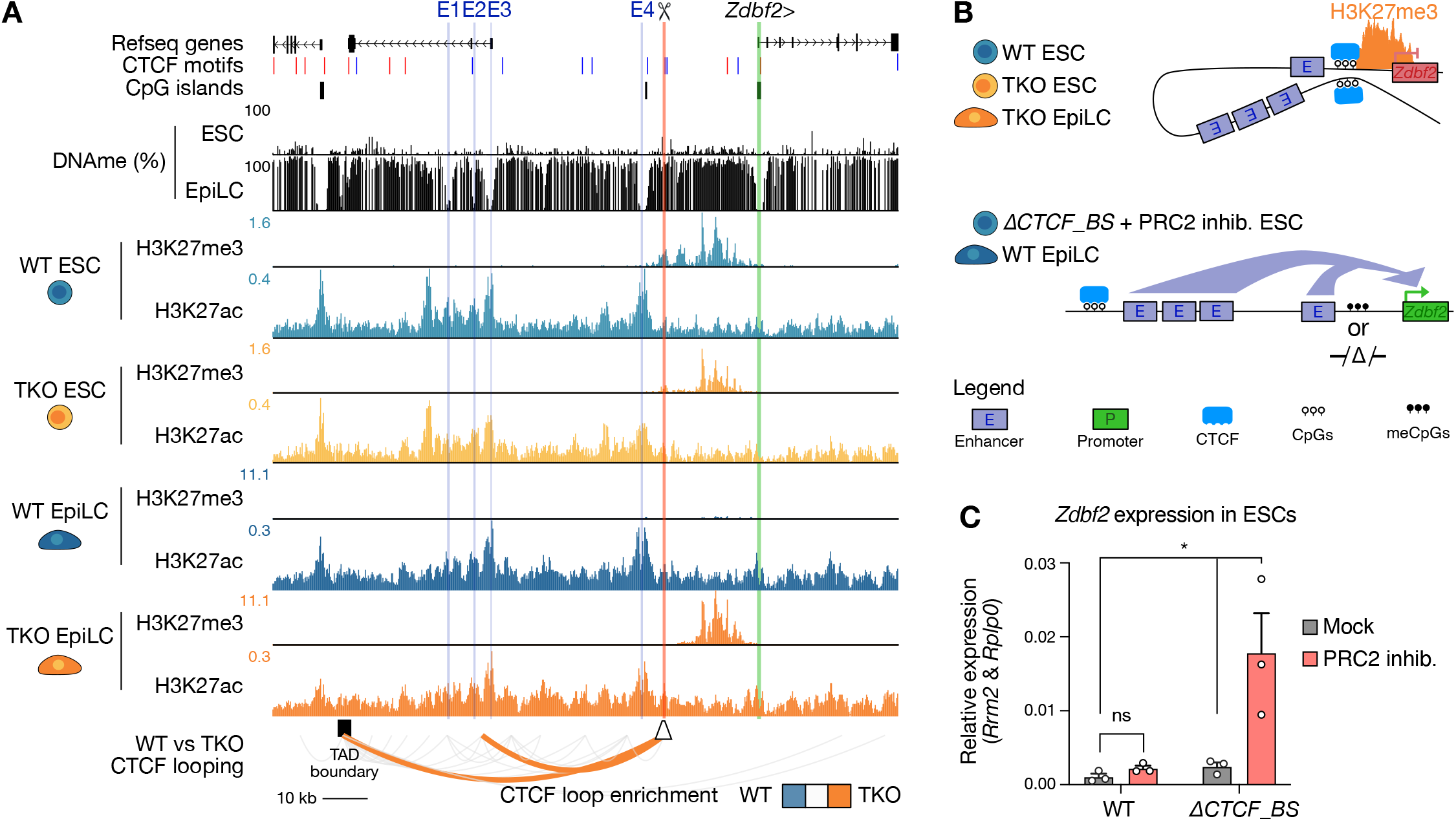
CTCF insulates promoter-enhancer interactions at the *Zdbf2* locus. **A.** UCSC genome browser screenshot of the *Zdbf2* locus. CpG methylation levels in WT ESCs and EpiLCs are shown, followed by H3K27me3 (Richard Albert et al. 2023) and H3K27ac (HiChIP) profiles in WT and TKO ESC and EpiLCs. Refseq genes, CpG islands, CTCF binding motifs and the TAD boundary are shown. Previously defined enhancers (E1-4) are shown in blue, the *Zdbf2* CGI promoter in green, and the CTCF binding site deletion is highlighted in red. Statistically enriched CTCF loops in TKO EpiLCs (as shown in Figure 3) are included (orange). *Zdbf2*: chr1:63,165,972-63,304,123 **B.** Schema showing the hypothesized insulating effect of CTCF binding on *Zdbf2* promoter-enhancer looping. Of note, in hypomethylated cells where CTCF is bound and enhancer sequences are insulated, the promoter sequence is marked by PRC2-associated H3K27me3. **C.** Bar chart showing the relative expression levels of *Zdbf2* by RT-qPCR in WT and *ΔCTCF_Binding Site* (*ΔCTCF_BS*) ESCs treated with UNC1999 (PRC2 inhibitor) or UNC2400 (mock). Data are shown as mean ± standard error for three replicates. p-values were calculated by one-tailed paired t-test assuming equal variance: *p<0.05

## Discussion

The CTCF binding landscape varies substantially between cell types, most likely related to its role in orchestrating the *cis*-regulatory interactions that help define cell identity. One potential regulator of CTCF localization is DNA methylation, as 5meC antagonizes CTCF in certain contexts. Proper CTCF/5meC localization is essential for regulation of some genomic imprints, and CTCF/5meC misregulation is associated with certain cancers (Flavahan et al. 2016, 2019). Thus, we set out to determine if DNA methylation plays an important role in shaping not only CTCF binding, but also the underlying gene expression program during cellular differentiation. We utilized an ESC to EpiLC differentiation system as this particular trajectory is associated with a dramatic increase in global DNA methylation levels, thus we reasoned that a large proportion of potential CTCF binding sites might be impacted. Furthermore, it is possible to carry out this differentiation protocol in the absence of all DNA methylation machinery; hence, we could focus specifically on the elements where CTCF remains enriched in the DNA methylation mutant. Crucially, we implemented epigenome editing to directly test the role of the methyl-mark *per se*, on CTCF binding and gene regulation.

Given the dramatic genome-wide gain of 5meC, one could conclude that only 1% of CTCF binding sites impacted represents a minor effect. On the other hand, 1% comprises about 1,000 sites, and impaired CTCF binding could directly influence the expression of hundreds or thousands of genes. This upper limit would constitute a substantial effect on genome regulation. Indeed, at day four of EpiLC differentiation, 1,459 genes were misregulated in TKO cells (FC≥2, adjusted p ≤ 0.05). We therefore felt it was necessary to distinguish CTCF-dependent from CTCF-independent modes of gene regulation in the 5meC mutant. To do so, we implemented a chromosome conformation technique that would allow us to enrich for CTCF binding sites. By assessing the CTCF-CTCF loops enriched at potentially DNA methylation sensitive binding sites, we could interrogate gene expression differences between WT and TKO at adjacent genes. Only a limited number (15) exhibited significant expression change (FC≥2, adjusted p≤0.05). Thus, our analyses would suggest that a small fraction of the total number of misregulated genes was likely due to direct CTCF-mediated control. It should be emphasized that expression of a greater subset of genes could be impacted via downstream effects, and there were more subtle changes in looping and/or expression that did not meet our thresholds.

We went on to examine four loci due to the likely effect that 5meC had on CTCF binding and looping. We did not discover a clear biological process linking these four genes. *Csf1* encodes a macrophage-stimulating factor that is widely expressed across mouse tissues, and is important for maintenance of tissue-specific macrophage populations (Sehgal et al. 2021). *Mob3b* is part of the highly conserved monopolar spindle-one-binder (MOB) gene family, and while the protein products have no known enzymatic function, they are thought to be scaffold proteins, and are linked with a number of human diseases (Gundogdu and Hergovich 2019). *Nrp2* codes for a transmembrane protein that contributes to a number of signaling pathways that contribute to the cytoskeleton, angiogenesis, and cancer progression (Grandclement et al. 2011). Conversely, we have extensively described *Zdbf2*. Interestingly, although it is expressed during early embryonic stages, *Zdbf2* mutants exhibit no obvious phenotypes either in ESCs nor in the *in vivo* embryo; rather, *Zdbf2* expression appears important in the postnatal hypothalamus (Greenberg et al. 2017; Glaser et al. 2022). Thus, it is possible that while the genes we described in this study were protected from ectopic CTCF-mediated gene control in WT EpiLCs (and potentially the *in vivo* epiblast) via 5meC deposition, we uncovered regulatory mechanisms that are biologically relevant in other cell types at later developmental stages. One could imagine that modulating the DNA methylation at the CTCF binding sites—via natural TET-mediated demethylation, for example (Wiehle et al. 2019), akin to what we did with our artificial dCas9 system—could tune the expression of the linked genes to ensure proper cellular function.

Related to this point, a substantial number of potentially DNA methylation sensitive binding sites could be masked due to the fact that even in WT EpiLCS, when global 5meC levels were elevated, the CTCF-bound region remained hypomethylated (Figure 1E). CTCF itself may protect from *de novo* DNA methylation at a substantial number of its binding sites (Lienert et al. 2011; Pant et al. 2003; Kemp et al. 2014). Additionally, these elements are likely protected by TET protein activity (Wiehle et al. 2019) and/or combinatorial transcription factor binding (Lienert et al. 2011; Krebs et al. 2014; Kremsky and Corces 2020). In other words, there is potential that DNA methylation could be substantially disruptive to the 3D regulatory structure of the genome absent of factors that deter DNA methylation machinery. This is hinted at by the fact that in some cancers, where the DNA methylome is broadly misregulated, there are notable examples of aberrant 5meC accumulation leading to CTCF loss resulting in oncogene expression (Flavahan et al. 2016, 2019), or repression of a tumor suppressor (Rodriguez et al. 2010). A previous study examined CTCF binding in a mouse ESC line harboring mutations in *Tet1* and *Tet2* (as opposed to our study here, the cells were cultured in conditions such that DNA methylation levels were globally high) (Wiehle et al. 2019). They observed that increased DNA methylation and loss of CTCF binding adjacent to promoters led to reduced gene expression; however, they employed a metastable cellular state and did not formally assess the impact of misregulated CTCF on chromatin conformation. Fodder for future studies will be to remove protection mechanisms, such as TET enzymes or pertinent transcription factors, in order to observe if ectopic gain of DNA methylation at discrete sites will have a more severe impact on CTCF-regulated genes than a complete loss of 5meC.

We utilized the EpiLC differentiation technique because it allowed for us to interrogate a cellular transition in the complete absence of DNA methylation. In most differentiated cell types DNA methylation is absolutely required, and acute 5meC loss leads to widespread epigenetic misregulation and cell death (Jackson-Grusby et al. 2001). However, EpiLCs are distinguished from other highly DNA methylated cell types in that they are still pluripotent, and express many of the same transcription factors that are linked with the pluripotency network (Takahashi et al. 2018). It is possible that this property is directly related with the resilience EpiLCs show in the absence of DNA methylation. In other words, there could be the same mechanisms in place in EpiLCs that allow DNA hypomethylated naïve ESCs to proliferate without fatal genomic instability. This could include ensuring that CTCF-mediated genome architecture largely stays intact independently of the underlying DNA methylation state. It is possible that in other more differentiated cell types, acute depletion of DNA methylation may lead to a more drastic effect than we observed in our TKO EpiLCs.

How then to bypass the cell death phenotype in DNA methylation mutants? Many chromatin conformation studies take advantage of degron technology and assay genome folding in the window between protein depletion and cell death. Such techniques have been successfully utilized to understand the role of CTCF (Nora et al. 2017), cohesin (Rao et al. 2017; Wutz et al. 2017), the Mediator complex (El Khattabi et al. 2019), and RNA polymerase II (Jiang et al. 2020b; Sun et al. 2021; Zhang et al. 2023). With high resolution techniques, such as HiChIP or Micro-C (Krietenstein et al. 2020; Hsieh et al. 2015, 2020), a degron system can be coupled with an assessment of the *cis*-regulatory interactome (Zhang et al. 2023). Such techniques could be adapted for DNA methylation degrons (eg, DNMT1) in differentiated cell types in order to gauge the impact of 5meC on the 3D genome.

Nevertheless, our EpiLC system did reveal a number of DNA methylation-sensitive CTCF binding events. The emergence of epigenome editing has enabled the direct assessment of the effect of a chromatin modification at a locus of interest without generating genetic mutants that exhibit potential confounding effects. Not only are these powerful tools in cell culture systems, as we described here, but they can also be implemented *in vivo* (Liu et al. 2016). Indeed, a SunTag/TET system highly similar to the one we utilized here has been successfully employed in mouse embryos to target the *H19-Igf2* imprint, which disrupted CTCF binding and *Igf2* expression (Horii et al. 2020, 2022). The prospect of using epigenome editing in the developing embryo proper to modify chromatin architecture and assess the physiological consequences presents a compelling endeavor for future studies.

## Materials and Methods

### ESC cell lines

E14Tg2a (E14) mouse ESCs was the parental line used for all experiments in this study, as well as serving as the background for all transgenic lines. The TKO was previously generated in-house (Dubois et al. 2022).

### Cell culture and differentiation

For the cells grown in serum culture conditions we used Glasgow medium (Gibco) supplemented with 15% Fetal bovine Serum (FBS), 0.1 mM MEM non-essential amino acid, 1mM Sodium Pyruvate, 2mM L-Glutamine, Penicillin, Streptomycin, 0.1 mM β-mercaptoethanol and 1000 U/ml Leukemia Inhibitory Factor (LIF). To pass, the cells were washed with 1X PBS, then trypsin was added to detach and disaggregate the cells for 5 minutes at 37°C. The desired number of cells were then transferred to a new flask.

For the 2i+vitC culture conditions we used N2B27 medium (50% neurobasal medium, 50% DMEM) supplemented with N2 (Gibco), B27 (Gibco), 2 mM L-Glutamine, 0,1 mM β-mercaptoethanol, Penicillin, Streptomycin, LIF and 2i (3 μM Gsk3 inhibitor CT-99021, 1 μM MEK inhibitor PD0325901) and Vitamin C (Sigma) at a final concentration of 100 μg/ml. To pass the cells, the media was removed, then Accutase (Gibco) was added to detach and disaggregate the cells and incubated for 5 minutes at room temperature. The desired number of cells were then transferred to a new plate. The ESCs in both conditions were grown on 0.1% gelatin-coated flasks in an incubator at 37°C and 5% CO2.

To induce EpiLC differentiation, cells were gently washed with PBS, dissociated, and replated at a density of 2 × 10^5^ cells/cm^2^ on Fibronectin (10 μg/ml, Sigma)-coated plates in N2B27 medium supplemented with 12 ng/ml FGF2 (R&D) and 20 ng/ml Activin A (R&D). EpiLCs were passed with Accutase at D3 of differentiation when the differentiation time was 7 days.

To inhibit PRC2 activity, cells were treated with 2 μM UNC1999 (Tocris) for at least one week. Control cells were treated with UNC2400 (Tocris)— an analog with >1,000 fold lower potency—at the same concentration.

### Generation of sgRNA constructs for epigenome editing

A piggyBac transposition compatible vector (Zuin et al. 2022) was modified by removing the sequences between the inverted terminal repeats by restriction digest, and incorporating a Hygromycin B resistance gene and U6-TRACR sequence by Gibson assembly. For *Nrp2*, a one guide RNA sequence was inserted by digesting the vector with BbsI, and ligating a double stranded DNA sequence containing compatible overgangs. For the other targets, dual-guide constructs were generated by linearizing the plasmid with BbsI and inserting a PCR product containing one gRNA sequence, the invariant sgRNA scaffold sequence, a modified murine U6 promoter and a second gRNA sequence, using the pLKO.1-blast-U6-sgRNA-BfuA1-stuffer plasmid as a template for amplification (Holoch et al. 2021). Guide RNA sequences were designed using the CRISPOR online program (crispor.tefor.net). Oligo sequences can be found in Supplementary Table S2.

### Generation of transgenic ESCs

All transgenesis experiments were performed with ESCs cultured in serum-containing media. Briefly, in each transfection ∼5 million cells were transfected with a mix containing 2,5 μg of each plasmid and plated at different concentrations to allow clone selection. We then performed electroporation using the Amaxa® Nucleofector® II Device from Lonza with the mouse ESC (A-013) program according to the manufacturer’s instructions. The transfected cells were cultured for a day in antibiotic-free media, and then were placed under antibiotic selection. The SunTag/TET epigenome editing construct was obtained from Addgene (Plasmid #82559). Individual clones that were Geneticin® (ThermoFisher) resistant were screened for Cas9 and GFP expression by Western blotting. The dual guide vectors were co-transfected with a plasmid containing the PiggyBac transposase, and Hygromycin B (ThermoFisher) resistant cells were pooled.

### Generation of CTCF binding site deletion

The deletion of the CTCF binding site was generated by transfecting two CRISPR sgRNAs flanking the target sequence along with Cas9. sgRNAs were designed using the online CRISPOR online program (crispor.tefor.net) and cloned into the pX459 plasmid harboring the Cas9 gene. Around five million WT serum-grown ESCs were transfected with 1μg of plasmids using Amaxa 4d Nucleofector (Lonza) and plated at a low density. Ninety-six individual clones were picked and screened by PCR for ∼600 bp deletion. Mutated alleles were confirmed by Sanger sequencing of cloned PCR amplicons. sgRNA sequences and genotyping primers can be found in Supplementary Table S2.

### CUT&RUN

We performed CUT&RUN according to the original protocol (Skene and Henikoff 2017) with the following modifications: 500,000 cells were used for each sample, and the primary antibody was incubated overnight at 4°C on a rotator. After incubation with pAG-MNase and performing the MNAse reaction, the samples were placed on a magnetic rack and the supernatant containing the DNA samples was recovered. Following addition of 0.1% SDS and 0.17 mg/ml Proteinase K, samples were incubated at 50°C for 1h. Purified DNA was obtained by phenol/chloroform extraction and precipitated with 100% ethanol by centrifugation. The DNA pellet was washed in 80% ethanol, spun down and air-dried before being resuspended in 25 μl of 1mM Tris-HCl pH 8.0. We used a primary CTCF antibody (Cell D31H2) or IgG control (SigmaAldrich I5006) and no secondary antibody was used for these experiments. The pAG-MNase plasmid was obtained from Addgene (#123461), and the protein was purified by the Curiecoretech Recombinant Protein Platform.

Sequencing library preparation was made using the NEBNext® UltraTM II DNA Library Prep Kit for Illumina (NEB) following the procedure described in “Library Prep for CUT&RUN with NEBNext® UltraTM II DNA Library Prep Kit for Illumina® (E7645) V.1” available on protocols.io. Quality control for the finalized libraries was performed using a TapeStation 420 system (Agilent). Libraries were sequenced by Novogene Co on a NovaSeq using paired-end 150 base pair parameters, requesting 4 gigabytes of data per sample, or approximately 13 million reads. The full list of datasets generated in this study are listed in Supplementary Table S1.

### Hi-ChIP

Hi-ChIP experiments were performed using the Arima Hi-ChIP kit (Arima Genomics) according to the manufacturer’s instructions. 15 μg of chromatin were used per sample and experiments were performed in duplicates. Briefly, cells were cross-linked with 2% formaldehyde for 10 min at room temperature, lysed and chromatin was digested with two different restriction enzymes included in the kit. Overhangs were filled-in in the presence of biotinylated nucleotides, followed by ligation. Ligated DNA was sonicated using the Covaris M220 to an average fragment size of 500 bp with the following parameters (Peak incident power: 50; Duty factor: 10%; Cycles per burst: 200; Treatment time: 250s). DNA was then immunoprecipitated overnight using 2.5 μg of H3K27Ac (Active Motif 91193) or CTCF antibody (Active Motif 91285). After a double-size selection to retain DNA fragments between 200 and 600 bp using Ampure XP beads (Beckman Coulter) the biotin-ligated DNA was precipitated with streptavidin-coupled magnetic beads (included in the kit).

The Hi-ChIP libraries were prepared on beads using the Accel-NGS 2D library Kit (Swift bioscience) following instructions from the Arima Hi-ChIP kit. Final libraries were analyzed using 4200 TapeStation system (Agilent) and sequenced.

### ChIP-qPCR

CTCF ChIP was performed as previously described (Schmidl et al. 2015). Briefly, 4 μl of CTCF rabbit antibody (AbFlex 91285) or 4 μl of IgG control rabbit antibody at 1 mg/ml (SigmaAldrich I5006) were combined to 50 μl of protein A magnetic beads (Invitrogen 10001D) and added to sonicated chromatin (from 200 to 700 bp, checked on agarose gel) from 7-9 million cells, O/N in the cold room. Beads were washed twice with TF-WBI (20mM Tris-HCl/pH 7.4, 150mM NaCl, 0.1% SDS, 1% Triton X − 100, 2mM EDTA), twice with TFWBIII (250mM LiCl, 1% Triton X-100, 0.7% DOC, and 10mM Tris-HCl, 1mM EDTA), and twice with with TET (0.2% Tween − 20, 10mM Tris-HCl/pH 8.0, 1mM EDTA). Chromatin was eluted and de-crosslinked in 70 μl of elution buffer (0.5% SDS, 300mM NaCl, 5 mM EDTA, 10mM Tris-HCl pH 8.0) containing 40 μg of proteinase K in an overnight incubation at 65°C. Eluted and purified DNA (Qiagen 28204) was directly used for qPCR. Data was first normalized to input, then to positive control locus (chr1:63181149-63181244). Primer sequences can be found in Supplementary Table S2.

### RNA extraction, cDNA synthesis and RT-qPCR

RNA extraction from cell pellets was performed using the KingFisher Duo Prime Magnetic Particle Processor and the MagMAX mirVana Total RNA kit, according to the manufacturer’s instructions.

First strand cDNA synthesis was performed using the SuperScript III Reverse Transcriptase kit (Invitrogen). 500 ng of total RNA were used for each reaction, along with 1 μL of 50 ng/μL of random primers, 1 μL of 10mM dNTP mix and sterile H_2_O up to 13μL. The rest of the procedure was performed following the manufacturer’s instruction.

For each RT-qPCR reaction, 1 μL of cDNA was mixed with 5 μL of LightCycler 480 SYBR Green I Master and 0.5 μL of 10 μM of each forward and reverse primers as well as sterile H_2_O up to 10 μL. The RT-qPCR was run on a LightCycler 480 II (Roche Applied Science) using 384 well plates. The samples first followed an initial incubation at 95°C for 10 minutes, and then 45 cycles of denaturation at 95°C for 10 seconds, annealing at 61°C for 20 seconds and extension at 72°C for 20 seconds. Samples were amplified in triplicates with appropriate non-template controls. Relative gene expression was calculated using the 2 −ΔCt method and normalized to the geometric mean of the expression levels of the two housekeeping genes *Rrm2* and *Rplp0.* Graphical representation and statistical analysis was performed with GraphPad Prism software. Primer sequences can be found in Supplementary Table S2.

### Protein extraction & Western Blot

For protein extraction, we used a BC250 lysis solution (25mM Tris pH 7.9, 0.2mM EDTA, 20% Glycerol, 0.25M KCl) supplemented with complete, EDTA-free protease inhibitors (Roche). Then, the samples were sonicated with a Bioruptor sonication device (High, 30 seconds on, 30 seconds off, for three cycles) and the protein concentrations were quantified using Pierce BCA Protein Assay Kit (ThermoFisher) on an Infiniate M200 (Tecan) machine. Western blot imaging was performed using the ChemiDoc MP (Biorad). The following antibodies and dilutions were used: Lamin-B1 (abcam ab16048) 1:2000 and H3K27me3 (Cell Signaling C36B11) 1:5000.

### Pyrosequencing

Genomic DNA was isolated from cells using the NucleoSpin Tissue kit (Macherey-Nagel). 500 ng-1 μg of genomic DNA was bisulfite converted using the EZ DNA Methylation-Gold kit (Zymo). Bisulfite-converted DNA was PCR amplified, and analyzed using the PyroMark Q24 machine and associated software (Qiagen). Graphical representation and statistical analysis was performed with GraphPad Prism software. Primer sequences can be found in Supplementary Table S2.

### LUMA

Genomic DNA (500 ng) was digested with MspI+EcoRI or HpaII+EcoRI (New England BioLabs) in parallel duplicate reactions. HpaII is a methylation-sensitive restriction enzyme, and MspI is its methylation-insensitive isoschizomer. EcoRI was included for internal normalization. The extent of the enzymatic digestions was quantified by pyrosequencing (PyroMark Q24), and global CpG methylation levels were then calculated from the HpaII/MspI normalized peak height ratio.

### WGBS analysis

Adapter and low-quality sequences were removed using Trimmomatic (v0.39) (Bolger et al. 2014) and parameters “ILLUMINACLIP:adapters.fa:2:30:10 SLIDINGWINDOW:4:20 MINLEN:24”. Read quality was assessed using FastQC before alignment to the mm10 genome using Bismark (v0.23.1) (Krueger and Andrews 2011) and default parameters. Reads with mates that did not survive read trimming or that could not be aligned in paired-end mode were concatenated and realigned. PCR duplicate reads were removed using deduplicate_bismark and CpG methylation information was extracted using bismark_methylation_extractor. Boxplots were generated in VisRseq (v0.9.42) (Younesy et al. 2015).

### CUT&RUN analysis

PE 150 reads were trimmed to 36 using Trimmomatic and parameter “CROP:36”. PCR duplicate reads were removed using Clumpify (v38.18) (Bushnell 2014) and parameters “dedupe=t k=19 passes=6 subs=$filter”, where “filter” is calculated by multiplying the rate of Illumina sequencing error (1%) with the read length. Subsequently, adapter-derived and low-quality nucleotides were removed using Trimmommatic as described above for WGBS. Read quality was assessed using FastQC before alignment to the mm10 genome using bowtie2 (v2.4.5) (Langmead and Salzberg 2012) and parameters “--local --very-sensitive --no-mixed --dovetail --no-discordant -- phred33 -I 10 -X 700”. Bigwig files were generated using Deeptools (v3.5.1) (Ramírez et al. 2016) bamCoverage and parameter “--normalizeUsing CPM --blackListFileName blacklisted_regions.fa -- ignoreForNormalization chrX chrM chrY --binSize 1”, removing blacklisted regions defined by the Kundaje lab. Aligned reads were used to call peaks using SEACR (v1.3) (Meers et al. 2019) and parameters “0.01 non stringent”. Peaks from all samples were merged using Bedtools (v2.30.0) (Quinlan and Hall 2010) and default parameters, resulting in 102,761 peaks. Genomic distribution of peaks was assessed using ChIPseeker (Wang et al. 2022). Enrichment of CTCF binding was calculated over peaks using VisRseq and RPKM values were used to calculate Spearman correlations using Morpheus (https://software.broadinstitute.org/morpheus). Venn diagrams were generated using pybedtools and matplotlib. PCA plots were generated using the sklearn PCA package (Pedregosa et al. 2012) and matplotlib. Average DNAme levels over CTCF peaks was calculated using Bedops (v2.4.40) (Neph et al. 2012). Differential CTCF enrichment over peaks was calculated using Limma (v3.54.1) (Ritchie et al. 2015) and default parameters. Scatterplots and violin plots were generated using VisRseq and matplotlib. FIMO (v5.5.0) (Bailey et al. 2009) was used to determine predicted CTCF binding motifs in the mm10 genome using the MA0139.1 downloaded from the 2022 JASPAR database (Castro-Mondragon et al. 2022).

### HiChIP analysis

HiChIP libraries were sequenced at shallow depth (3-4 million paired-end reads) and the ARIMA MAPS pipeline (v2.0) (Juric et al. 2019) was used to calculate target sequencing depth. Following deep sequencing, the HiC-Pro pipeline (v3.1.0) (Servant et al. 2015) was used to digest the mm10 genome (^GATC G^ANTC), align reads and generate contact map matrices. Single-end alignment bam files were used for peak calling using macs2 (v2.2.7.1) (Zhang et al. 2008) and generating bigwigs using bamCompare (as described above). Chromosome 11 consistently showed more reads in TKO cells and was omitted in subsequent analyses. Correlation between samples was assessed using HiCexplorer hicCorrelate over 5kb matrices. PCA plots were generated using fanc (v0.9.25) (Kruse et al. 2020) over chromosome 19 using parameters “-Z -s 100000”. A/B compartments were calculated using HiCexplorer hicPCA over 25kb matrices. CTCF enrichment over previously defined autosomal TAD borders (Bonev 2017) was caulculated using VisRseq. Loops were called using hichipper (v0.7.7) (Lareau and Aryee 2018b) and parameters “-mu -pp 1000” and “-mu -pp 1750” and differential loops were calculated using diffloop (Lareau and Aryee 2018a) and quickAssoc normalization. bigInteract tracks of differential looping were colour coded based on statistical significance (FDR<0.05). Virtual 4C plots were generated in HiCexplorer (v3.7.2) (Wolff et al. 2020, 2018; Ramírez et al. 2018) using a combination of chichViewpoint (-- averageContactBin 4 --range 500000 500000), chicSignificantInteractions (--pValue 0.2 -- xFoldBackground 2), chicAggregateStatistic (default parameters), chicDifferentialTest (--alpha 0.25 --statisticTest chi2) and chicPlotViewpoint (-- pValueSignificanceLevels 0.1). hicCompareMatrices and hicPlotMatrix was used to generate heatmaps. UCSC genome browser track hubs (Raney et al. 2014) were also generated for visualization.

### RNAseq analysis

PCR duplicate reads, as well as adapter and low quality sequences, were removed as described above. Trimmed reads were aligned to the mm10 genome using STAR (v2.7.9a) (Dobin et al. 2013) and default parameters. Gene expression levels were quantified over Refseq genes using VisR and uniquely aligned reads (MAPQ=255). Differential expression analysis was conducted using Limma and default parameters. Bigwigs were generated as described above with the additional parameter “--minMappingQuality 255”. Bar charts were generated using matplotlib (Hunter 2007).

## Availability of data and materials

High throughput sequencing data was uploaded to NCBI GEO under accession number GSE246984. See Supplementary Table S1 for the full list of data analyzed in this study. Custom scripts are available under an GNU General Public License v3.0 on GitHub: https://github.com/julienrichardalbert/3DNAmethylation/releases/tag/v0.0.

## Acknowledgments

We thank Roshani Singh and Julie Segueni for assistance with the project. We thank Ana Paula Azambuja for sharing differential loop-calling R scripts and Joël Marchand for maintaining computational resources. We thank members of the Greenberg lab for useful discussions. Work in the Greenberg group is supported by the European Research Council (ERC-StG-2019 DyNAmecs), a Laboratoire d’excellence Who Am I? (Labex 11-LABX-0071) Emerging Teams Grant and funds from the Agence National de Recherche (ANR, project ANR-21-CE12-0015-03). A.M.S. is supported by Fondation pour la Recherche Médicale (FRM) (SPF202004011789) and ARC (ARCPDF12020070002563) postdoctoral fellowships. J.R.A is supported by a FRM, Post doc France Fellowship (SPF202110014238). Work in the Noordermeer group is supported by funds from the ANR (projects ANR-21-CE12-0034-01, ANR-22-CE12-0016-03 and ANR-22-CE14-0021-02) and PlanCancer (19CS145-00).

## Author Contributions

A.M-S., J.R.A. and M.V.C.G. conceived the study and designed the experiments. A.M-S. and M.S. performed the experiments. J.R.A. performed the genomic analyses, with assistance from A.M-S and D.N.. D.N. provided guidance for HiChIP analyses and interpretation. A.M.S., J.R.A. and M.V.C.G. wrote the paper. All authors edited the paper.

---

**Supplementary Figure S1.**
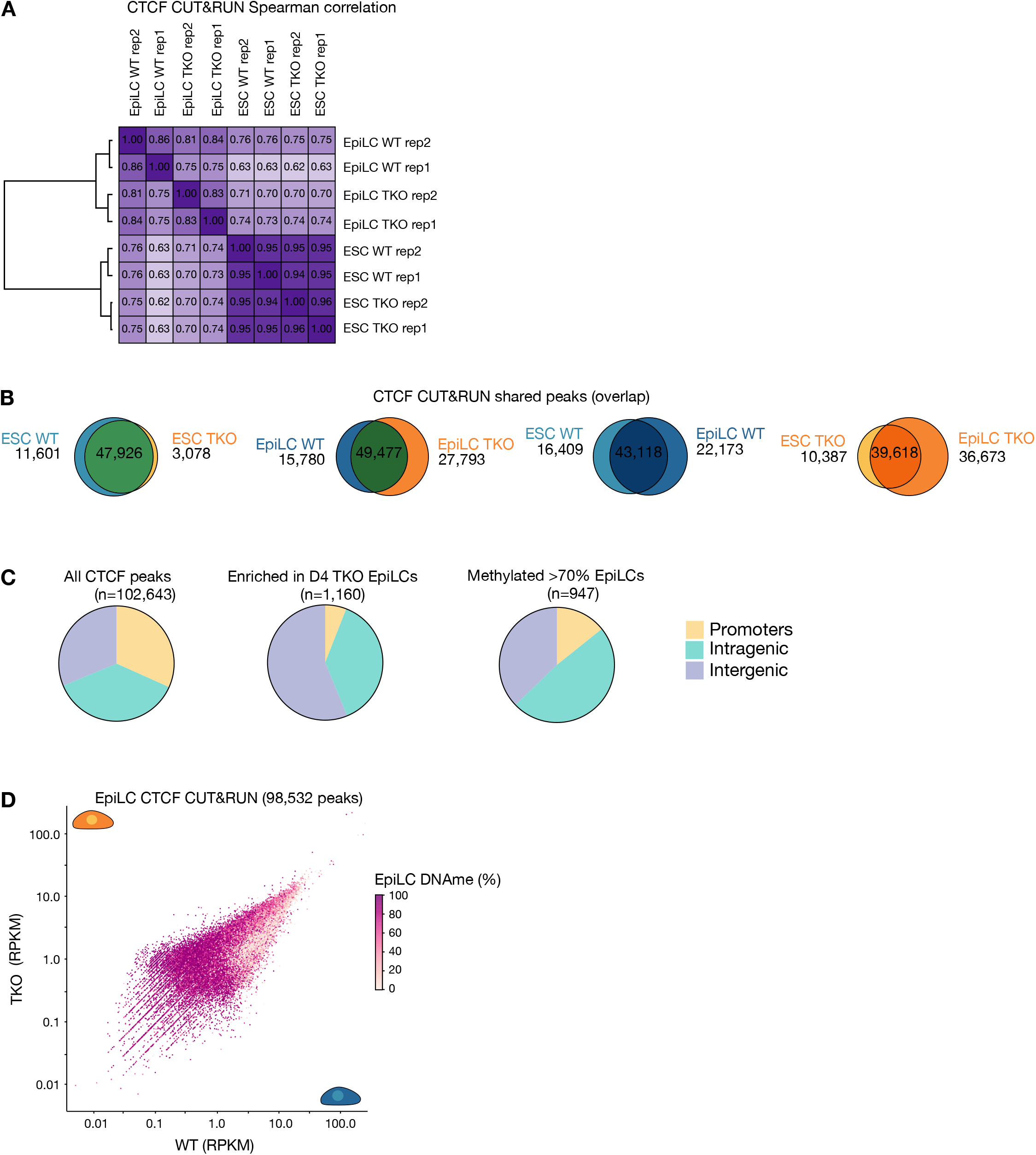
Profiling CTCF occupancy in WT and TKO ESCs and EpiLCs by CUT&RUN. **A.** Correlogram showing the Spearman correlation of CTCF enrichment over CTCF peaks in WT and TKO ESCs and EpiLCs. Hierarchical clustering of datasets is included (left). **B.** Overlap between CTCF peaks in WT and TKO ESCs and EpiLCs. The number of peaks unique to and overlapping between cells are indicated. **C.** Pie charts showing the distribution of CTCF binding sites over promoters, intragenic regions (gene bodies) and intergenic regions from the CUT&RUN data. CTCF binding sites are categorized based on CTCF enrichment in TKO EpiLCs (FC>2, adj. pval<0.05), and those enriched in TKO EpiLCs that normally gain >70% 5meC in WT EpiLCs. **D.** 2D scatterplot showing CTCF enrichment over CTCF peaks in WT and TKO EpiLCs as in Figure 1. Data points are colored based on mean 5meC levels in WT EpiLC. Only peaks with at least one CpG for which 5meC levels could be assessed (5X read coverage) are shown (n=98,532).

**Supplementary Figure S2.**
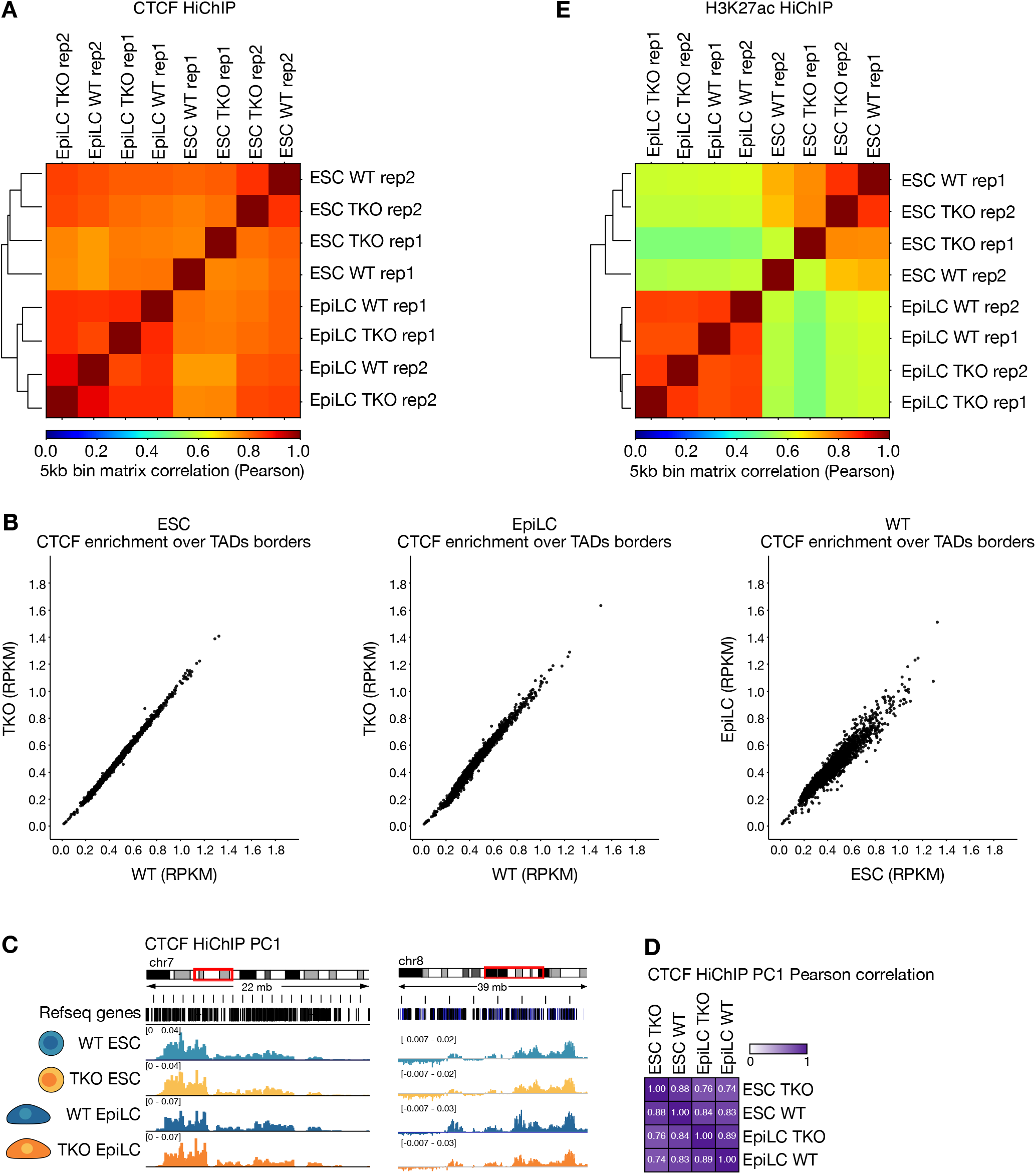
Profiling 3D CTCF interactions in WT and TKO ESCs and EpiLCs. **A.** Correlogram depicting the Pearson correlation of 3D contacts over 5kb bins from CTCF HiChIP in WT and TKO ESCs and EpiLCs. **B.** 2D scatter plots showing CTCF enrichment (RPKM) over TAD borders in WT and TKO ESCs and EpiLCs (n=2,426). **C.** Integrated Genome Browser screenshots showing TAD compartmentalization (PC1) using CTCF HiChIP data. **D.** Correlogram showing Pearson correlation levels of TAD compartmentalization (PC1) over 100 kb bins using CTCF HiChIP data. **E.** Correlogram showing the Pearson correlation of 3D contacts over 5kb bins from H3K27ac HiChIP in WT and TKO ESCs and EpiLCs.

**Supplementary Figure S3.**
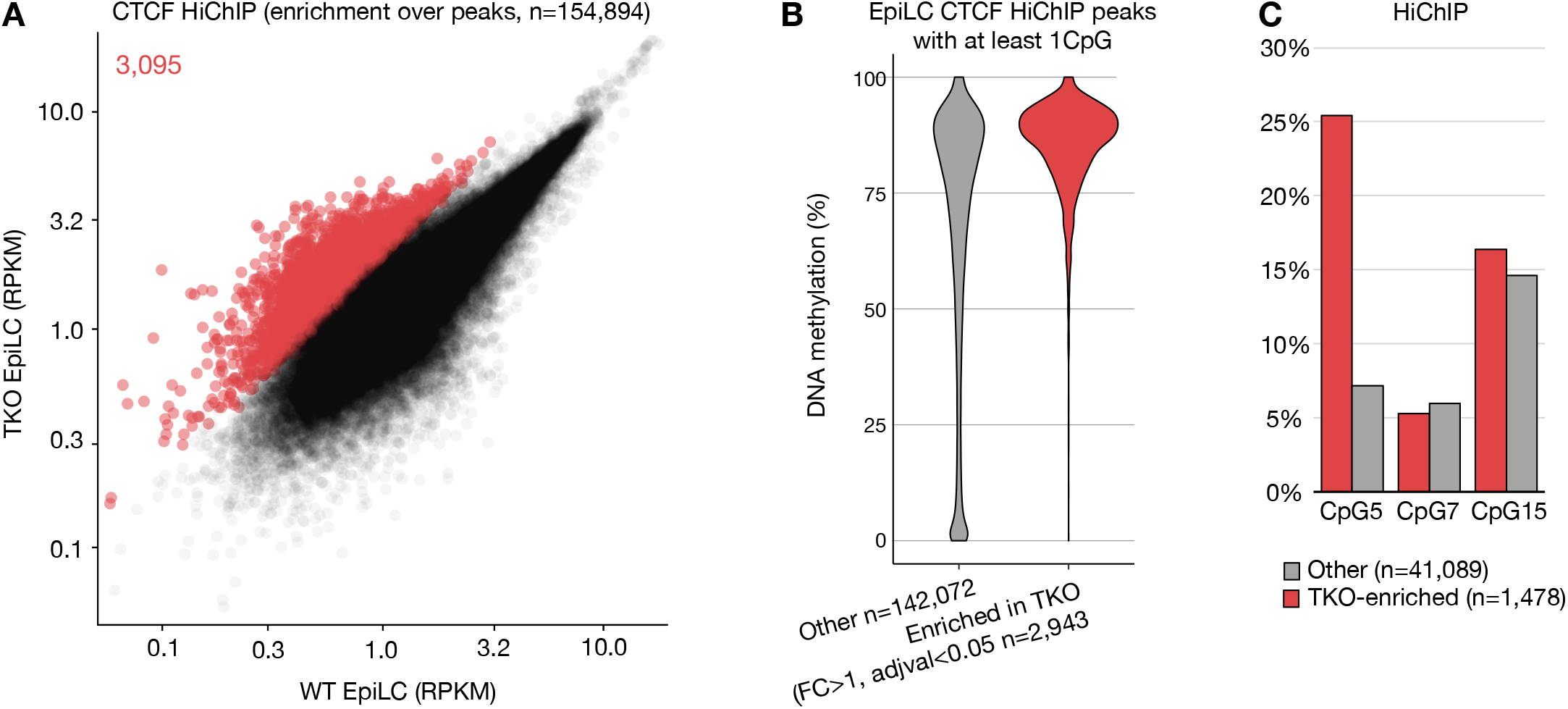
CTCF exhibits potential 5meC sensitivity in CTCF HiChIP data. **A.** 2D scatterplot showing CTCF enrichment (RPKM) over CTCF peaks in WT versus TKO EpiLC HiChIP data. CTCF peaks enriched in TKO EpiLCs are highlighted in red. **B.** Violin plot of the distribution of EpiLC CpG methylation levels within CTCF peaks. Peaks are categorized as in A. **C.** Bar plot showing the proportion of CTCF peaks that overlap a canonical CTCF binding motif with a CpG at position 5, 7 or 15.

**Supplementary Figure S4.**
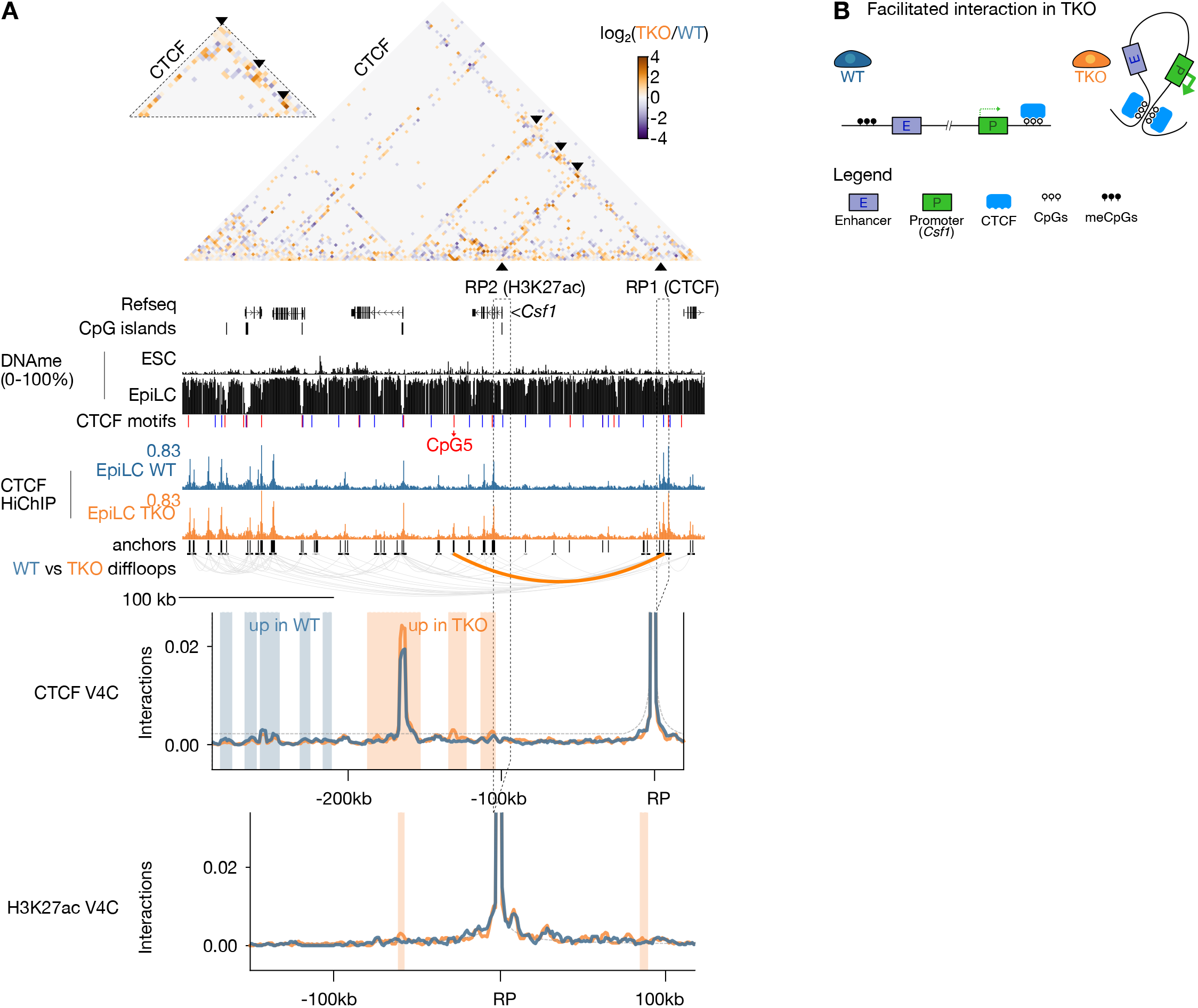
5meC-sensitive CTCF looping is associated with increased promoter-enhancer contacts and gene expression at the *Csf1* locus in TKO EpiLCs. **A.** Contact matrix (top), genome browser screenshot (middle) and virtual 4C (bottom) plots of the *Csf1* locus. Top: differential CTCF contacts between WT and TKO EpiLCs are displayed, where each pixel represents a 1 kb bin. The inset magnifies the TKO-enriched CTCF contacts. Reference points (RP) 1 (CTCF) and 2 (H3K27ac) for the virtual 4C plots (bottom) are indicated by a dashed rectangle. Middle: browser screenshot showing CpG methylation, CTCF enrichment levels and TKO-enriched CTCF loops. Refseq genes, CpG islands, CTCF motifs (positive strand in red, negative strand in blue) are included. The CTCF motif with a CpG at position 5 that underlies a putative 5meC-sensitive CTCF peak is indicated in red. Bottom: Virtual 4C plots of reference points 1 (CTCF) and 2 (H3K27ac) showing interaction frequencies between the reference point and adjacent area. The background model is shown as a dotted gray line. Statistically enriched contacts (chi-squared test, alpha<0.25) are highlighted in blue (high in WT EpiLC) or orange (high in TKO EpiLC). Coordinates: chr3:107,554,568-107,890,568. **B.** Schema depicting the hypothesized facilitated enhancer-promoter interactions and increased gene expression in TKO EpiLCs.

**Supplementary Figure S5.**
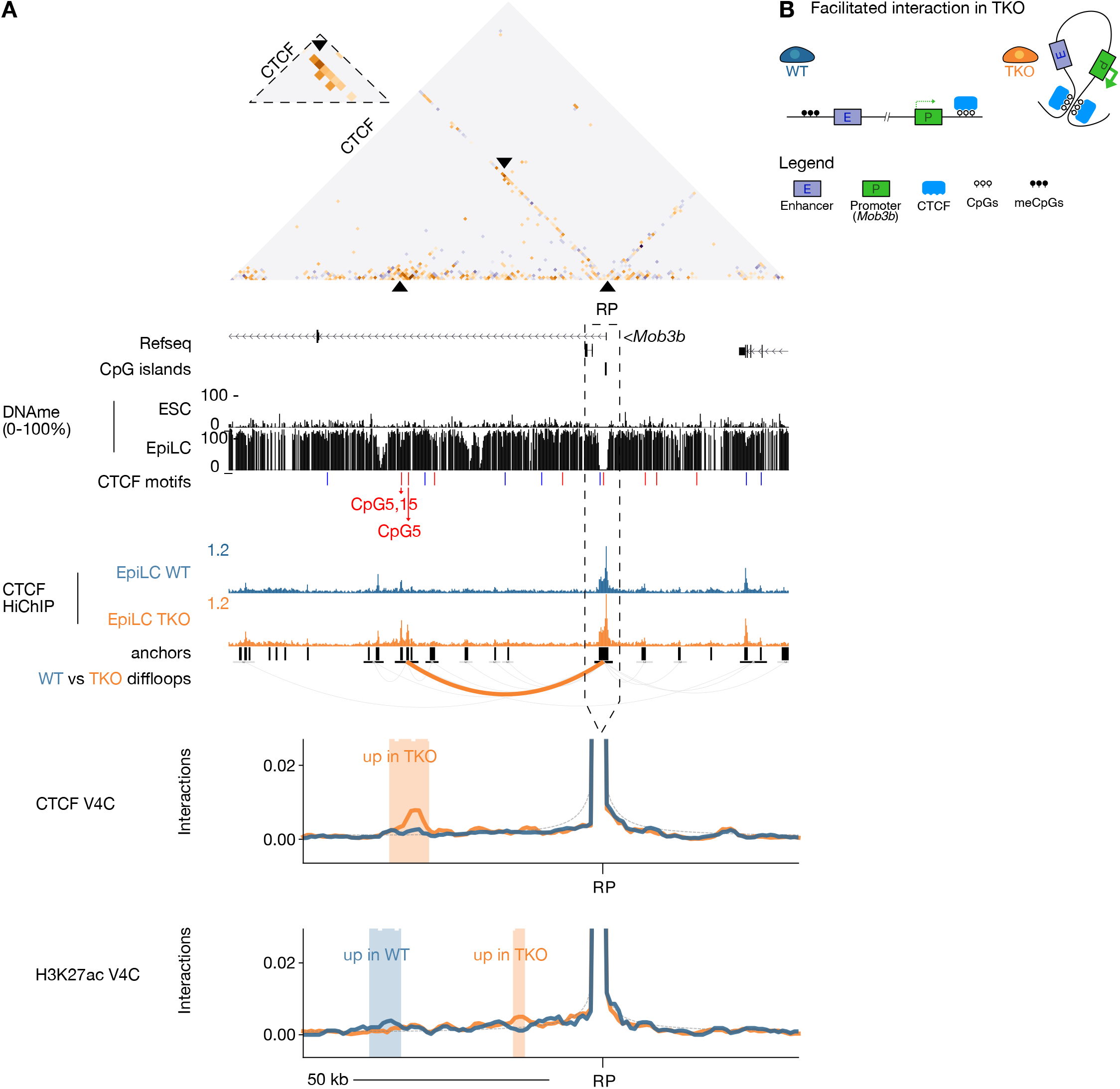
5meC-sensitive CTCF looping is associated with increased promoter-enhancer contacts and gene expression at the *Mob3b* locus. **A.** Contact matrix (top), genome browser screenshot (middle) and virtual 4C (bottom) plots of the *Mob3b* locus as in Supplementary Figure S4. Coordinates: chr4:35,061,303-35,203,887. **B.** Schema depicting the hypothesized facilitated enhancer-promoter interactions and increased gene expression in TKO EpiLCs.

**Supplementary Figure S6.**
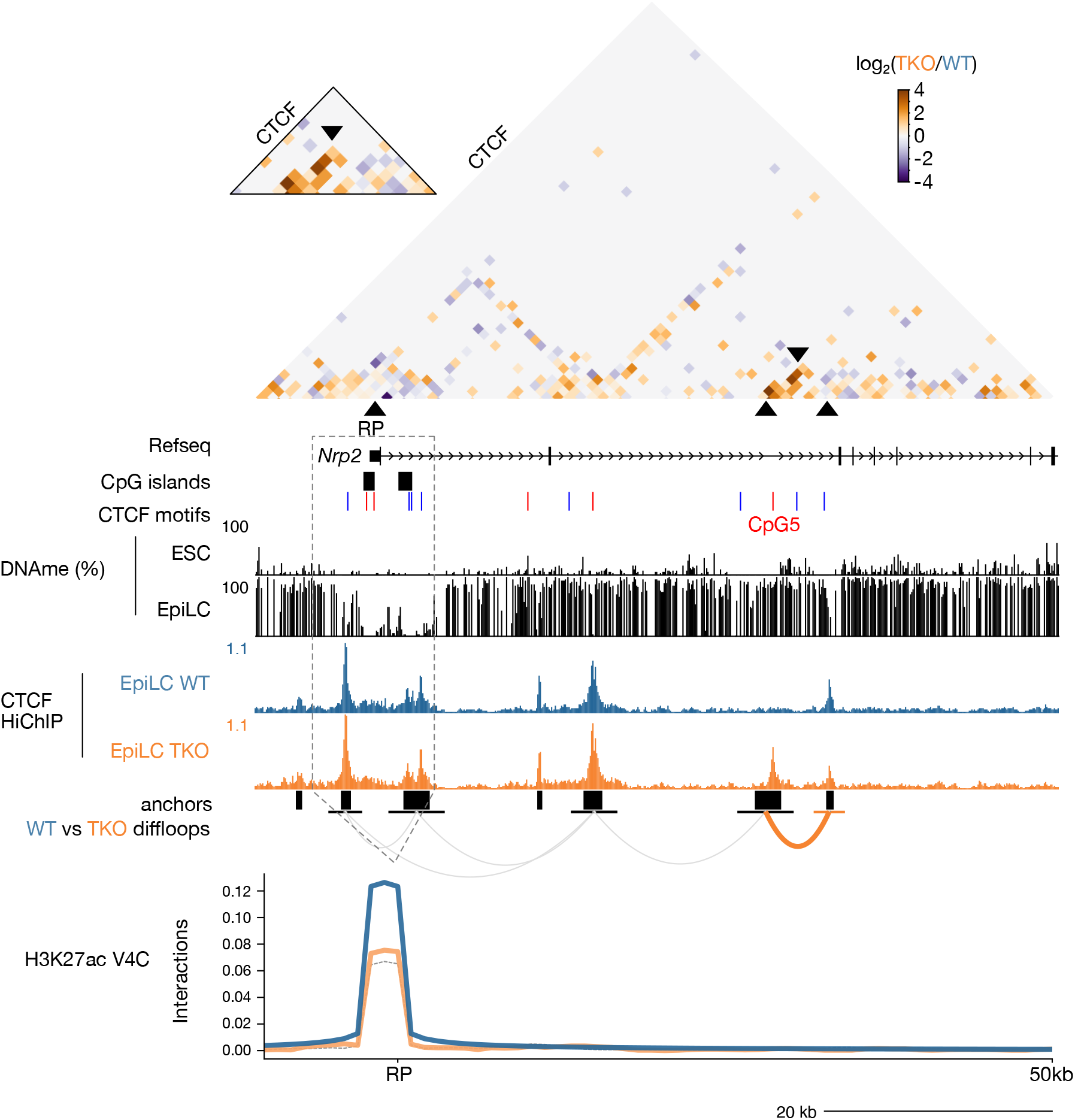
**5meC-sensitive CTCF looping at the *Nrp2* locus.** Contact matrix (top), genome browser screenshot (middle) and virtual 4C (bottom) plots of the *Nrp2* locus as in Supplementary Figure S4. Note the lack of differential H3K27ac interactions between the promoter and adjacent sequences. Coordinates: chr1:62,693,316-62,763,316.

**Supplementary Figure S7.**
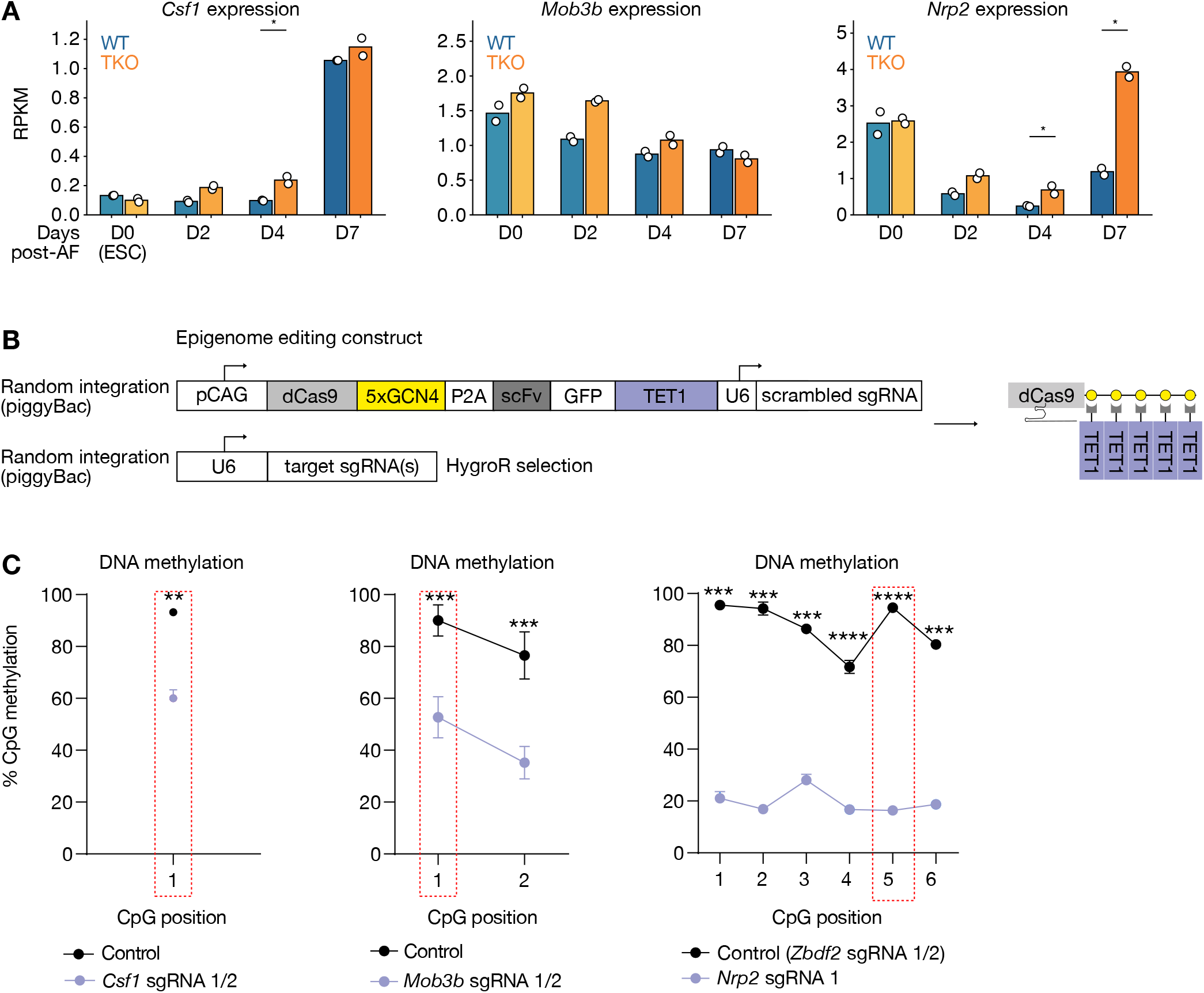
Site-directed cytosine demethylation at candidate 5meC-sensitive CTCF loops. **A.** Heatmap showing expression levels (RPKM) of genes near 5meC-sensitive CTCF loops over a 7-day EpiLC differentiation time course. Time points where genes are scored as differentially expressed (linear modeling using Limma, t-test, log_2_FC>1, adj. p value<0.05) are indicated by an asterisk. **B.** Epigenome editing construct. A constitutive CAG promoter drives expression of a catalytically inactive Cas9 (dCas9) fused to 5xGCN4 epitopes (SunTag), a self-cleavable peptide (P2A), and a single chain variable fragment (scFv) that recognizes GCN4 epitopes fused to GFP and the human TET1 catalytic domain. This construct also contains a scrambled sgRNA sequence under control of a U6 promoter. A separate construct containing U6 promoter driving expression of targeted sgRNAs was transfected for experiments and individual loci. Both constructs were stably inserted in the genome by piggyBac transposition; the dCas9-SunTag/TET1 construct was selected by sorting GFP expressing cells, and the sgRNA construct by hygromycin selection. **C.** Bisulfite-pyrosequencing results of epigenome edited cell lines. The CpG corresponding to CpG5 in the canonical CTCF binding motif are highlighted by red boxes. Data are shown as mean ± standard error for three replicates. p-values were calculated by one-tailed paired t-test assuming equal variance: *p<0.05, **p<0.01, **p<0.001, ****p<0.0001.

**Supplementary Figure S8.**
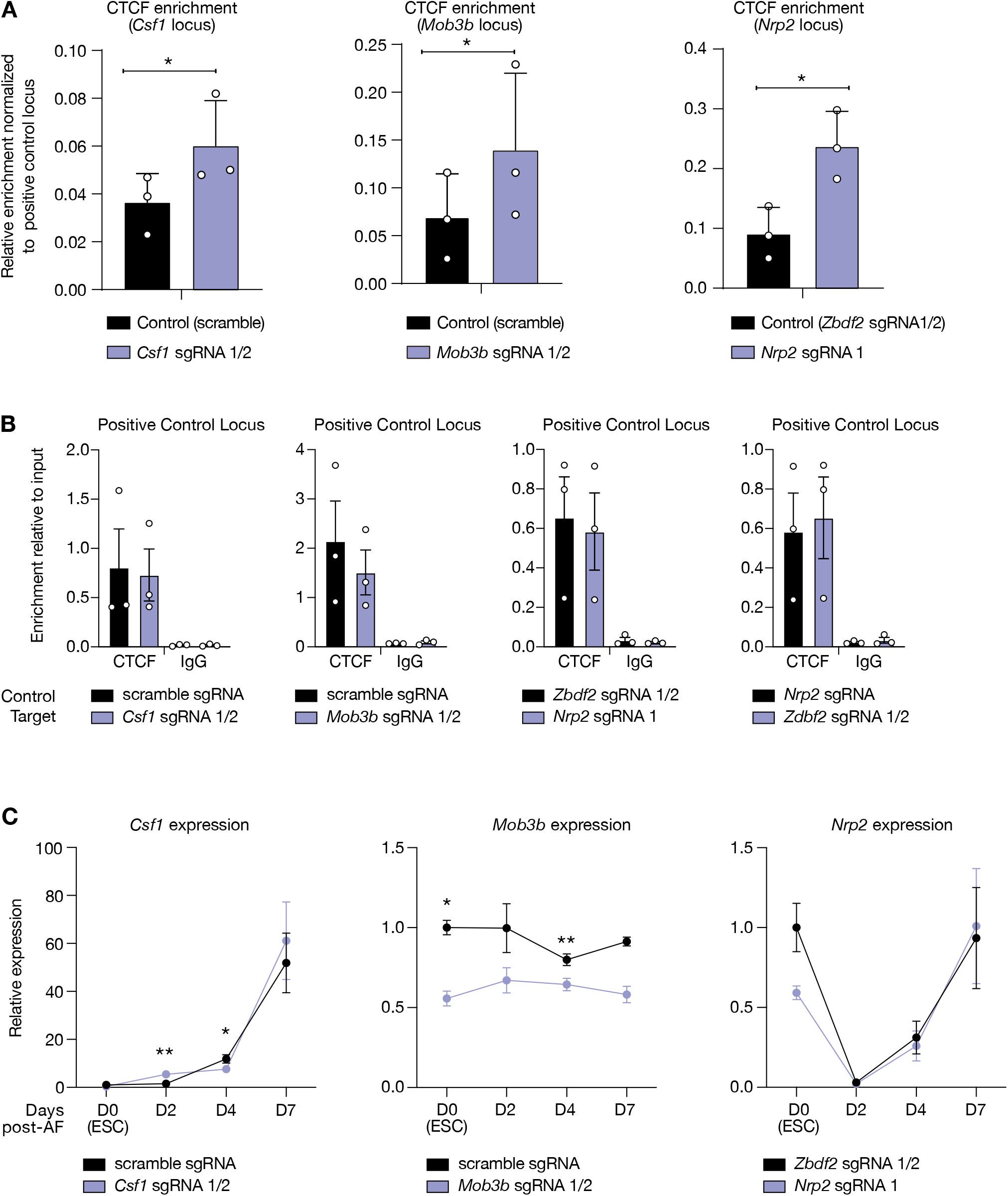
Site-specific cytosine demethylation results in increased CTCF recruitment but modest changes in nearby gene expression levels. **A.** CTCF ChIP results with targeted cytosine demethylation. Data are shown as mean ± standard error for three replicates represented by unfilled circles. **B.** CTCF and IgG (background) ChIP at positive control locus (chr1:63181149-63181244) relative to input DNA. Data are shown as mean ± standard error for three replicates represented by unfilled circles. **C.** RT-qPCR results of the same cells as in A over a time course of 7 days of EpiLC differentiation. Expression of each replicate was normalized to two housekeeping genes (*Rrm2* & *Rplp0*), and then to WT ESCs. Data are shown as mean ± standard error for three replicates. Data are shown as mean ± standard error for three replicates. p-values were calculated by one-tailed paired t-test assuming equal variance: *p<0.05, **p<0.01, **p<0.001, ****p<0.0001.

**Supplementary Figure S9.**
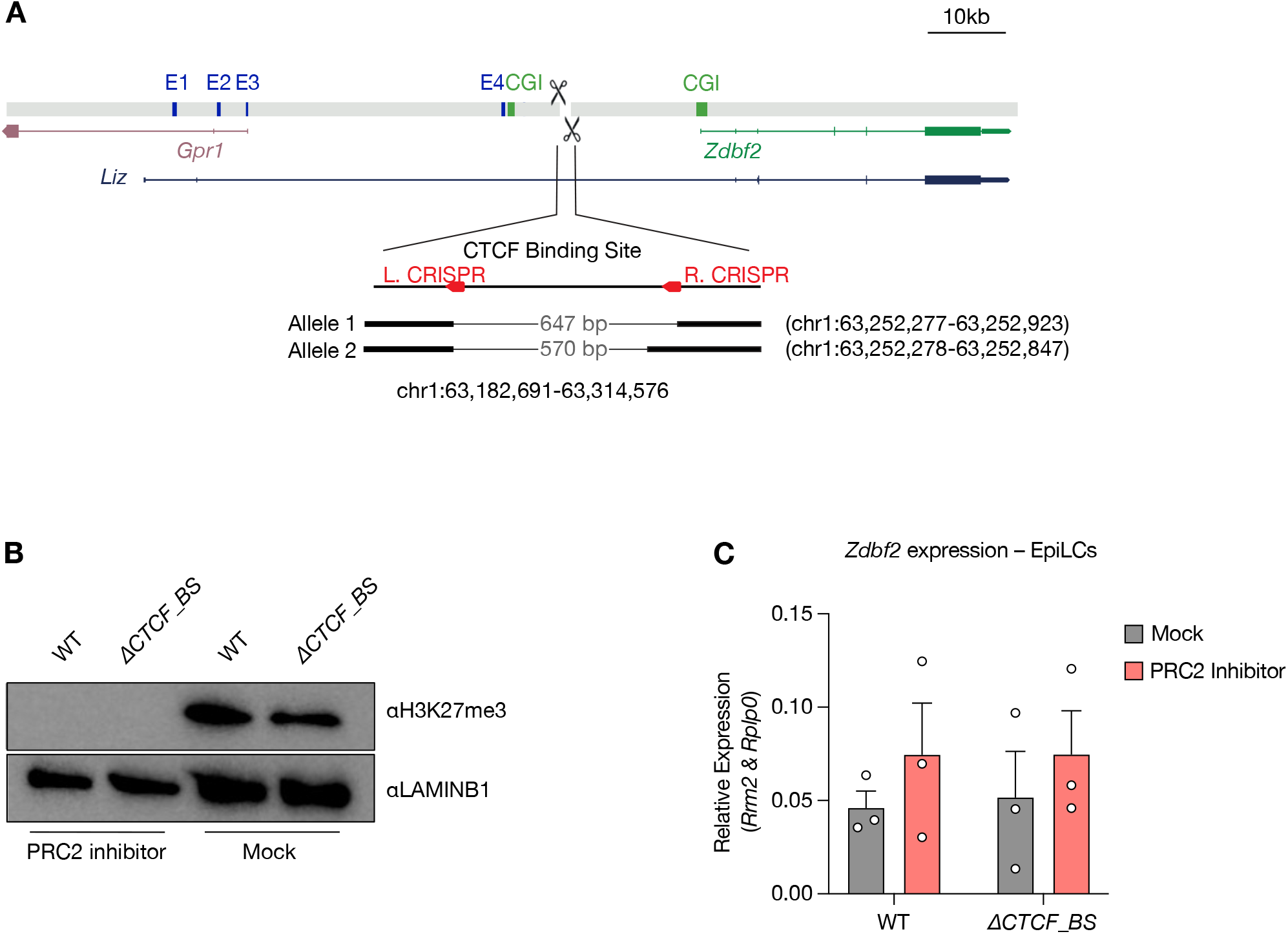
Genetic ablation of a 5meC-sensitive CTCF binding site. **A.** Schema depicting target Cas9 target sites (red), previously defined enhancers (E1-4, blue) and CpG islands (CGI, green) at the *Zdbf2* locus. The entire window spans chr1:63,182,691-63,314,576 and the length and coordinates of the deletions are indicated. **B.** Western blot showing H3K27me3 levels in WT and CTCF binding site-deleted (*ΔCTCF_BS*) ESCs grown in 2i+vitC. Cells were treated with UNC1999 (PRC2 inhibitor) or UNC2400 (mock). LAMINB1 was used as a loading control. **C.** Relative *Zdbf2* expression levels in EpiLCs treated with UNC1999 or UNC2400. Expression levels were normalized to housekeeping genes *Rrm2* and *Rplp0*. Data are shown as mean ± standard error for three replicates, and individual replicates.

## Notes

### Competing Interest Statement

The authors have declared no competing interest.

